# Non-redundant cardiolipin synthase family proteins shape lipid composition and stress resilience in *Bacteroides fragilis*

**DOI:** 10.1101/2025.05.12.653583

**Authors:** Matthew K. Schnizlein, Bongjin Hong, Jennifer N.T. Nguyen, Katarina Jones, Alyssa I. Rodriguez, Aretha Fiebig, Shawn R. Campagna, Marcy J. Balunas, Thomas V. O’Halloran, Sean Crosson

## Abstract

Gut-resident bacteria must withstand membrane-disrupting stresses, such as bile acids, to persist in the intestinal environment. The Bacteroidota are a dominant phylum of the human gut microbiota. However, the genes responsible for the synthesis of membrane lipids in this phylum remain relatively uncharacterized. *Bacteroides fragilis* is a prevalent member of this phylum and encodes two predicted cardiolipin synthase family proteins, ClsA and ClsB. We previously identified both *cls* genes as bile-acid fitness factors in *B. fragilis* P207, but the contributions of *clsA* and *clsB* to cell physiology remained unclear. Here, we combine targeted gene deletion with high-resolution lipidomics, metabolomics, and elemental mass spectrometry to show that ClsA and ClsB have distinct and non-redundant functions in the cell. Each *cls* gene differs in its growth-phase expression pattern, and deletion of each gene produces distinct effects on cell morphology, fitness under membrane-perturbing stress, and the broader metabolome. *clsA* and *clsB* strongly influence the levels of different glycyl-seryl peptidolipids, a *Bacteroides* lipid family that includes immune-active lipids, and whose loss impairs bile acid resistance and gut colonization. In contrast to the acute ion-gradient disruption caused by the secondary bile acid deoxycholate, deletion of *clsA* and *clsB* did not measurably alter steady-state intracellular ion levels under standard growth conditions. Together, these results define related but non-redundant roles for two cardiolipin synthase family proteins in a common member of the human gut microbiota.

**Importance:** The gut bacterium *Bacteroides fragilis* is a normal member of the human intestinal microbiota that can also bloom to high abundance in inflamed guts. To survive within the gut, *B. fragilis* must maintain the integrity and function of its cell membrane. In this study, we report that the two cardiolipin synthase family genes in the *B. fragilis* genome impact the cell’s lipid profile in distinct ways and contribute differently to how *B. fragilis* responds to membrane stress. Our work provides insight into how a common gut bacterium adapts to conditions encountered in the intestine and improves understanding of membrane biology in this important group of gut microbes.

## Introduction

To colonize gut niches and persist within the host, bacteria must maintain membrane integrity against host immune pressures and membrane-disrupting molecules such as bile acids (1–3). The functional properties of membranes are shaped by the lipid classes they contain and by structural variation within those classes (4–6). For example, bacteria can remodel phospholipid acyl chains in response to extracellular conditions, altering membrane fluidity and/or permeability through changes in both chain length and unsaturation. They may also alter membrane properties by changing phospholipid abundance (4, 6) or by modifying lipid headgroups, including through aminoacylation reactions that can stabilize bilayers (7, 8). Since membrane lipids are drawn from shared biosynthetic pools, altering the synthesis of particular lipid classes can also have consequences that extend beyond the membrane to the broader metabolic economy of the cell.

A conserved class of enzymes that contributes to bacterial membrane lipids is the cardiolipin synthase (Cls) family. Most bacterial Cls proteins belong to the phospholipase D superfamily and contain two conserved HKD motifs that together form a catalytic center (9). Their phosphatidyl-donor and acceptor specificities vary (10, 11), and some members can yield products other than cardiolipin (10, 12, 13). Bacteria commonly encode multiple Cls-family proteins (14, 15), raising the question of whether these enzymes are functionally redundant or have evolved distinct roles. This question has been addressed in only a few organisms and remains open across most bacterial lineages. It is especially pertinent in the Bacteroidota, a dominant phylum of the human gut microbiota whose putative cardiolipin synthase family proteins are largely uncharacterized.

*Bacteroides fragilis*, which encodes two Cls-family proteins, is a good model for addressing this knowledge gap in the phylum Bacteroidota. It is an anaerobic, gram-negative gut resident (16) that can use nanomolar oxygen as a terminal electron acceptor (17) and is clinically relevant as both a beneficial symbiont and an opportunistic pathogen (18). Enterotoxigenic strains encoding *B. fragilis* toxin/fragilysin have been linked to intestinal disease and colorectal cancer (19), while non-toxigenic *B. fragilis* is common in the healthy gut, where HMP-based analyses estimate a mean stool relative abundance of ∼1.5% (20). Its fitness in the gut depends in part on cell-envelope functions that support membrane integrity (21, 22), including membrane lipid biosynthesis (23). Sphingolipids, including diverse ceramide species, are prominent membrane constituents in *Bacteroides* (24, 25). The *B. fragilis* group additionally produces glycine-, serine-, and glycyl-seryl-containing lipids, including families in which an acylated glycyl-serine moiety is linked through phosphate to a diacylglycerol or a dihydroceramide backbone (24). We recently identified two cardiolipin synthase family genes (*clsA* and *clsB*) as important for the fitness of the non-toxigenic *B. fragilis* strain P207 under exposure to 0.01% deoxycholate (21), a secondary, microbially modified bile acid; this concentration falls within the range measured in the healthy human gut (26). *B. fragilis* P207 was originally isolated from the ileoanal pouch of a patient with ulcerative colitis (20) and is representative of *B. fragilis* populations that can dominate prior to the onset of pouchitis, a condition characterized by inflammation of the ileoanal pouch following colectomy (27).

In this study, we investigate the roles of the two *B. fragilis* cardiolipin synthase family (*cls*) genes. *B. fragilis* P207 ClsA and ClsB fall into two conserved genus-level phylogenetic groups. Deletion of either gene perturbs subsets of membrane lipids, including distinct effects on the glycyl-seryl conjugates of diacylglycerol (glycyl-seryl phosphatidic acid [GS-PA]) and of dihydroceramide (glycyl-seryl phospho-dihydroceramide [GS-PDHC]), and has separable effects on fitness during stress and on cell morphology. Given known links between membrane composition and maintenance of cellular ion gradients, we experimentally defined the baseline ion content of *B. fragilis* and tested whether *cls* gene loss perturbs intracellular ion homeostasis, finding that steady-state ion levels are maintained under standard conditions despite the envelope stress phenotypes of the mutants. Loss of each gene also produces separable changes in the broader metabolite pools. We conclude that *clsA* and *clsB* make distinct contributions to cellular lipid composition, cell morphology, and stress resilience.

## Results

### *B. fragilis* P207 ClsA and ClsB make distinct contributions to membrane lipids

The two *B. fragilis* P207 genes, ptos_000612 (*clsA*) and ptos_003252 (*clsB*), encode proteins with predicted cardiolipin synthase domains (TIGR04265, PFAM13091, and cd09112) (Fig. 1A). Despite this shared domain architecture, ClsA and ClsB are only 29.6% identical to each other, and each is ∼30% identical to *Escherichia coli* ClsA (Fig. 1A). Low absolute identity is expected across these evolutionary distances: 11 conserved housekeeping proteins (AtpD, DnaK, FusA, GroL, GyrA, GyrB, RecA, RpoB, RpoC, SecA, and Tuf) share a median pairwise identity of only ∼55% across *B. fragilis* P207, *E. coli* K-12, and *B. subtilis* 168 (range ∼35%–76%). The cross-species Cls comparisons sit at or just below the low end of that spread, so the divergence is substantial but not significantly greater than that of proteins whose function is conserved. Sequence identity at this level, therefore, cannot resolve whether ClsA and ClsB catalyze the same reaction or different ones.

**Figure 1.**
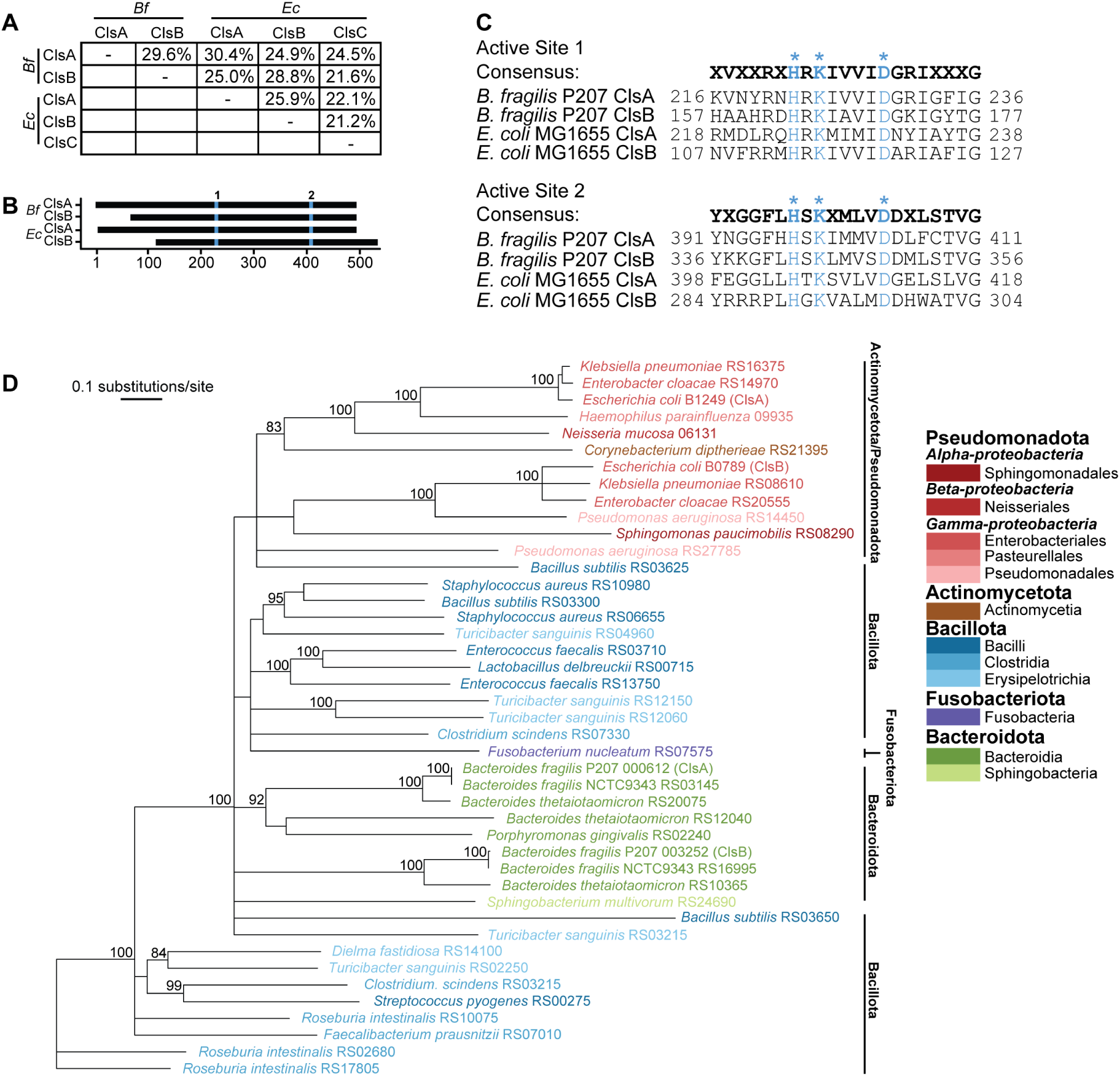
Structural analysis of *B. fragilis* P207 ClsA and ClsB, two cardiolipin synthase family proteins. (A) Pairwise amino acid sequence identity among cardiolipin synthases from *B. fragilis* P207 (Bf) and *E. coli* K12 MG1655. (B) Schematic alignment of ClsA and ClsB from *B. fragilis* P207 and *E. coli* MG1655, with the two HKD domains highlighted in blue. (C) Consensus and individual sequences of the regions flanking the two HKD active sites of *B. fragilis* and *E. coli* Cls proteins. Positions of the two HKD motifs are indicated in blue and marked with asterisks; the N- and C-terminal positions are shown. (D) Neighbor-joining phylogeny of Cls proteins from representative gut bacterial taxa, with one genome per species. Taxon names and locus tag numbers are shown at branch tips; full metadata are provided in Table S2. Bootstrap support values >80% are shown; 100 independent replicates were performed. Branch colors indicate phylum: red, Pseudomonadota (three classes; one, one, and three orders represented); brown, Actinomycetota (one class); blue, Bacillota (three classes); violet, Fusobacteriota (one class); and green, Bacteroidota (two classes). Scale bar indicates amino acid substitutions per site.

Sequence divergence between ClsA and ClsB is not, however, reflected at the structural level. AlphaFold3 (28) models of the two *B. fragilis* proteins predict highly similar global folds and catalytic cores, with a structural overlay yielding a root mean square deviation of 0.93 Å across 320 residues (Fig. S1B). Sequence conservation between *B. fragilis* and *E. coli* ClsA and ClsB is concentrated in two regions flanking the predicted active sites, with limited identity across the rest of the proteins (Fig. 1B and C). Both regions harbor a phospholipase-D-type H-X-K-X(4)-D motif (29, 30), consistent with phospholipase-D catalytic activity (Fig. 1C). *E. coli* ClsC, by contrast, has a divergent active site 1 sequence relative to *B. fragilis* ClsA/ClsB (Fig. 1B and C) and was therefore excluded from further comparisons.

To test the role of *clsA* and *clsB* in *B. fragilis* lipid synthesis, we deleted each gene individually and in combination and conducted lipidomic analysis on the strains. Each deletion resulted in the loss of distinct membrane lipids. MS-DIAL analysis (31) of the liquid chromatography-tandem mass spectrometry (LC-MS/MS) data identified minor features whose accurate masses are consistent with cardiolipin (Table S3), though no MS2 spectrum supports this annotation. These features were undetectable in both backgrounds lacking *clsB* and were not reduced in Δ*clsA*. We observed much stronger shifts in another family of lipids. Glycyl-serine (GS) peptidolipids (annotated as NAGlySer species in LipidBlast *in silico* MS/MS libraries) are abundant in *Bacteroides* (24) and showed large changes upon *cls* deletion. *clsB* deletion reduced the glycyl-seryl phospho-dihydroceramides by at least 100-fold in every species relative to wild type (WT), with 5 of the 12 species falling below the limit of detection; deletion of *clsA* reduced the glycyl-seryl phosphatidic acid lipids 12- to 45-fold. The *clsA clsB* double mutant eliminated GS-PA entirely and reduced GS-PDHC to the same extent as deletion of *clsB* alone (Table S3). All six GS-PA species responded in the same direction in Δ*clsA*, and none of them required *clsB* (Table S3). Glycyl-seryl lipid (GS-lipid), the presumptive precursor, was higher in the backgrounds lacking *clsB* for four of its five species. Our data are consistent with a model in which ClsA and ClsB contribute to the synthesis of GS lipid conjugates. A more comprehensive analysis of how the loss of *clsA* and *clsB* affects the broader membrane lipid composition of the *B. fragilis* envelope is presented in sections below.

### *Bacteroides* cardiolipin synthase family proteins form two conserved genus-level groups

The lipid changes described above were measured in a single strain, so we next asked whether *Bacteroides* genomes generally encode two distinct *cls* genes, examining Cls protein sequences and gene neighborhoods across the genus. To evaluate the relatedness of *Bacteroides* Cls proteins to homologs from other gut-associated bacterial lineages, we aligned Cls protein sequences from representative gut-associated taxa and constructed a phylogenetic tree. In this targeted comparison, Cls homologs largely grouped by bacterial lineage, with Bacteroidota sequences forming a distinct cluster and Bacillota sequences separating into two clusters (Fig. 1D). As this analysis was limited to representative gut-associated taxa, we interpret the tree as a focused comparison of the sampled sequences rather than a comprehensive reconstruction of bacterial cardiolipin synthase family evolution. Within this sampling, the topology is consistent with lineage-associated diversification of Cls proteins and does not suggest recent widespread horizontal transfer among the sampled gut-associated phyla.

We next evaluated Cls protein sequences within the *Bacteroides* genus. We aligned 301 *Bacteroides* protein sequences containing the TIGR04265 domain (i.e., cardiolipin synthases). Most genomes encoded two cardiolipin synthase homologs that partitioned into two primary-sequence clusters (Fig. S1). The *Bacteroides thetaiotaomicron* genome encodes a third Cls homolog that is distinct from the two major *Bacteroides* Cls clusters (Fig. S1), suggesting an evolutionarily divergent paralog or a lineage-specific acquisition. We designate cluster 1 homologs as ClsA because they group with the *B. fragilis* ClsA characterized in this study and share N-terminal hydrophobic features with *E. coli* ClsA (32, 33). We designate cluster 2 homologs as ClsB because they group with the second broadly conserved *Bacteroides cls* gene represented by *B. fragilis clsB*. We left the third *B. thetaiotaomicron* cls homolog unnamed and did not further characterize it. Within the *Bacteroides* genus, the chromosomal gene neighborhoods around *clsA* (Fig. S2A) and *clsB* (Fig. S2B) are highly conserved; this conservation is lost outside the *Bacteroides*. The conserved *clsA* neighborhood includes genes encoding an AAA-family ATPase, a DUF3822-family protein, an RsmD-family RNA methyltransferase, an outer membrane β-barrel protein, AroB, hypothetical proteins, and a nearby tRNA-Pro. The conserved *clsB* neighborhood is distinct and includes genes annotated as an IgA peptidase M64-family protein, an Lrp/AsnC-family regulator, dihydrofolate reductase, thymidylate synthase, DUF5056- and DUF6249-domain proteins, an RNA polymerase sigma factor, and an ORF6N-domain protein. Two separately conserved gene neighborhoods, each associated with one sequence cluster, are consistent with the two *cls* genes having been maintained independently within the genus rather than arising by recent duplication.

### *clsA* and *clsB* deletion mutants have distinct growth phenotypes under membrane stress

Membrane lipid composition governs the permeability barrier and the capacity to maintain transmembrane ion gradients. Having found that loss of ClsA and ClsB produces distinct changes in *Bacteroides* lipid classes implicated in gut colonization and stress resilience (23, 34, 35), we asked whether the corresponding mutants also differ in their sensitivity to agents that challenge cell envelope functions. We assessed the growth of *B. fragilis cls* deletion mutants under conditions that perturb membrane integrity, osmotic balance, pH homeostasis, or transmembrane ion gradients. In the absence of treatment, all strains grew similarly (Fig. 2; Fig. S3). Treatment with membrane stressors revealed *cls-*dependent growth defects. Detergent stressors, deoxycholate or SDS, impaired the growth of the *cls* mutants relative to WT, with the Δ*clsA* Δ*clsB* double mutant showing the strongest defect in deoxycholate; reintroduction of either *clsA* or *clsB* partially restored growth in the double-mutant background (Fig. 2A and B; Fig. S3). Loss of *clsB* had a larger effect than loss of *clsA*, particularly during SDS treatment, where Δ*clsB* strains showed a pronounced delay in growth.

**Figure 2.**
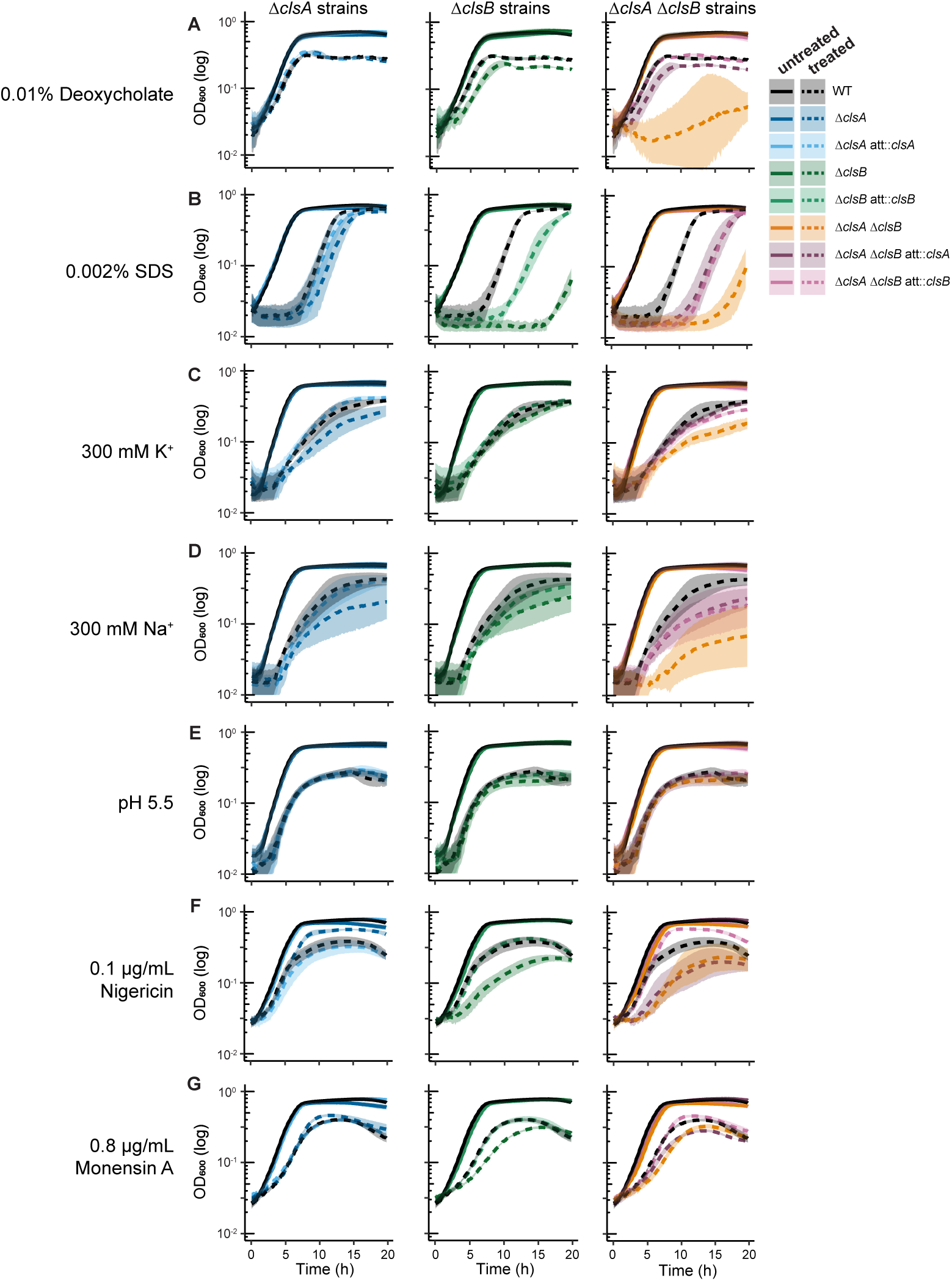
Loss of *cls* gene function results in growth defects under stress. Optical density (OD600) measurements of *B. fragilis* P207 WT and *cls* single or double mutant strains exposed to the detergents (A) 0.01% deoxycholate or (B) 0.002% SDS; the salts (C) 300 mM K+ or (D) 300 mM Na+; (E) medium adjusted to pH 5.5; and the ionophores (F) 0.1 µg/mL nigericin or (G) 0.8 µg/mL monensin A. Untreated (solid lines) and treated cultures (dashed lines) are plotted by strain, with blue, green, and orange/purple indicating *clsA*-, *clsB*-, and *clsA clsB*-related strains, respectively. Black indicates WT. The shaded areas around each line indicate two times the standard error of the mean of *n* = 3 biological replicates.

Elevated NaCl or KCl also revealed growth defects in the mutant panel, with the double mutant showing the most pronounced defect (Fig. 2C and D; Fig. S3). In contrast, lowering the medium pH impaired the growth of all strains without a clear *cls*-dependent effect (Fig. 2E). To test sensitivity to ion-gradient disruption more directly, we treated cells with nigericin, which primarily disrupts K+/H+ gradients, or monensin A, which preferentially disrupts Na+/H+ gradients (36). Under ionophore stress, strains lacking *clsB* were generally more impaired than WT (Fig. 2F and G; Fig. S3). Strains lacking *clsA* behaved in the opposite direction, reaching a higher maximum culture density than WT, an effect that was complemented by reintroduction of *clsA*. This difference met the threshold for statistical significance under monensin A, where Δ*clsA* also grew with a shorter doubling time than WT; the same trend was apparent under nigericin but did not reach significance (Fig. S3). Together, these growth phenotypes provide evidence that ClsA and ClsB make distinct contributions to *B. fragilis* membrane physiology, with ClsB playing a broader protective role across tested membrane-perturbing conditions.

### Deletion of *clsA* and *clsB* alters cell morphology

Strains lacking *clsB* lose the glycyl-seryl phospho-dihydroceramides, described as an abundant lipid family of the *Bacteroides* membrane (24); strains lacking *clsA* lose the less abundant glycyl-seryl phosphatidic acid lipids (Table S3). As envelope lipid composition contributes to the mechanical properties that set cell shape (37), we asked whether these compositional changes are reflected in *B. fragilis* cell morphology. Deletion of either *cls* gene did not markedly impact overall rod-like cell shape or cellular length distribution (Fig. S4A and B). However, strains lacking *clsA* showed a modest but significant reduction in cell width (i.e., Δ*clsA* and Δ*clsA* Δ*clsB* strains; Fig. 3A and B; Fig. S4C). Cells lacking *clsB* were not significantly narrower than WT, but this population showed a broader cell-width distribution, suggesting that ClsB contributes to cell-size homeostasis (*F*-test, WT to Δ*clsB P*= 0.019, WT to Δ*clsA* Δ*clsB* att::*clsA P* < 0.001; WT variances compared to Δ*clsA* and Δ*clsA* Δ*clsB* were not significantly different; Fig. 3A; Fig. S4C). Both phenotypes were genetically complemented. We conclude that loss of ClsA and ClsB activity produces modest effects on *B. fragilis* cell morphology (Fig. 3A and B).

**Figure 3.**
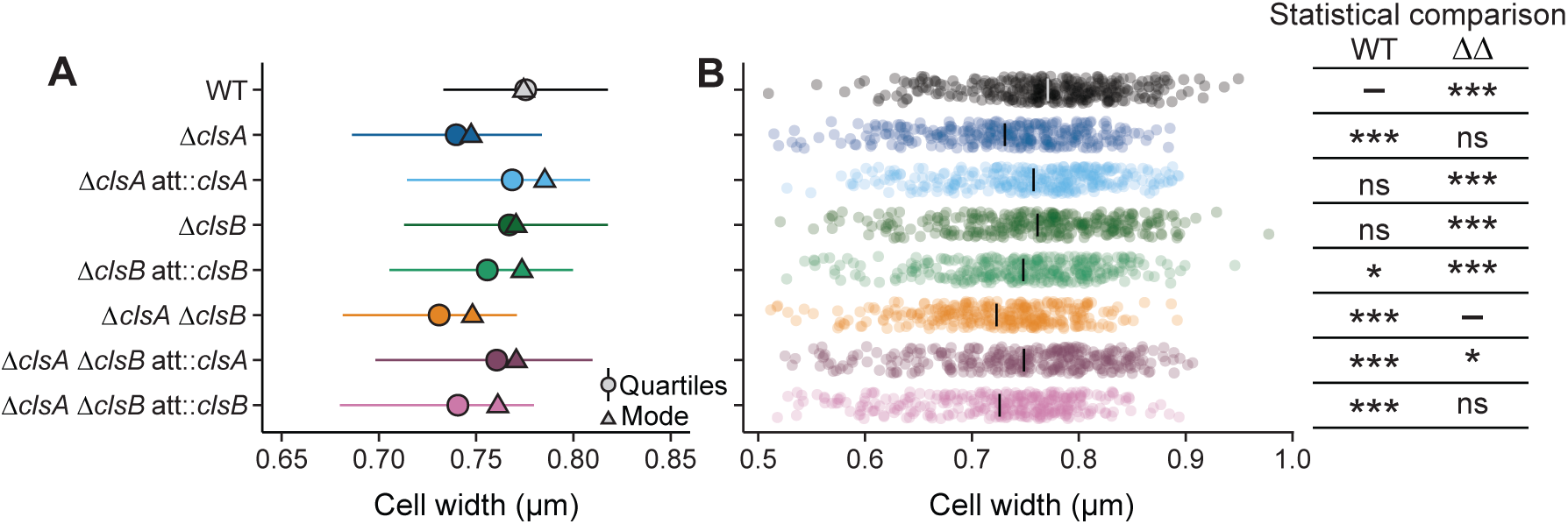
Loss of *cls* gene function is associated with defects in cell morphology. (A) Summary of the cell width distributions. Horizontal line lengths indicate the 25% and 75% quartiles of the distribution; circles indicate the median; triangles indicate the mode (i.e., the peak of the distribution). (B) Cell width data for all cells shown in panel A plotted by strain (*n* = 323 cells/strain). The gray or black bars indicate the mean. The table at right shows the results of a mixed-effect linear model comparing strain width to the widths of either WT cells or Δ*clsA* Δ*clsB* double mutant (ΔΔ) cells, as indicated at the top of the column, with a random intercept of biological replicate (i.e., cell width ∼ strain + 1 biological replicate). Unadjusted *P*-values are shown; *, *P* < 0.05; ***, *P* < 0.001.

### *B. fragilis cls* genes have distinct expression profiles

Since Δ*clsA* and Δ*clsB* mutants showed distinct stress phenotypes in *B. fragilis*, we asked whether the two genes also show different patterns of regulation. Prior RNA-seq analysis of WT *B. fragilis* P207 showed that both genes are moderately expressed relative to the transcriptome: *clsA* transcript levels were near the genome-wide median, while *clsB* levels fell above the median but below the upper quartile (21). We examined how growth phase affects the expression of each gene and found that *clsB* transcript levels were approximately fourfold higher during logarithmic phase than stationary phase, while *clsA* showed the opposite pattern, with approximately twofold higher expression in stationary phase (Fig. 4A). Thus, *clsA* and *clsB* have reciprocal growth-phase-dependent expression profiles.

**Figure 4.**
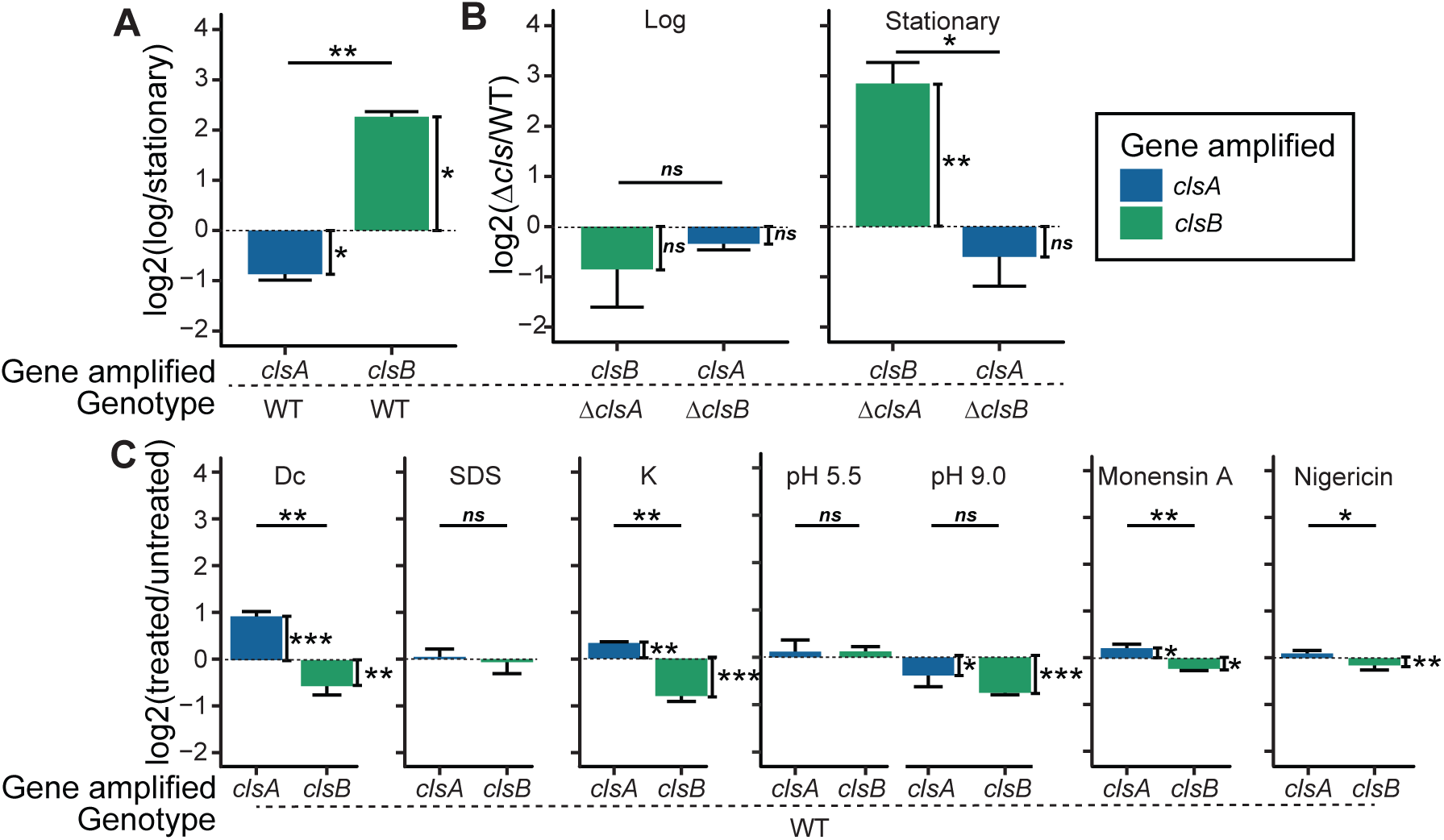
Cardiolipin synthase family genes have distinct expression profiles. (A) RT-qPCR analysis of relative levels of *clsA* and *clsB* transcripts, shown as the log2 ratios between log- and stationary-phase growth. (B) Expression levels of *clsA* or *clsB* when the other gene is knocked out in log- or stationary-phase growth. (C) *clsA* and *clsB* expression levels in WT *B. fragilis* P207 after 20-min exposure to 0.01% deoxycholate, 0.002% SDS, 300 mM K+, pH 5.5 and 9.0, 0.8 µg/mL monensin A, or 0.1 µg/mL nigericin. For all panels, values are means ± standard deviation from three biological replicates, each biological replicate value being the mean of three technical replicates. The vertical brackets for statistical analysis assess the significance of the difference between conditions in panels A and C and between mutant and WT in panel B. The horizontal brackets for statistical analysis assess the significance of the difference between ΔΔCt values for *clsA* and *clsB* (Student’s *t*-test). Genes amplified are colored in blue (*clsA*) or green (*clsB*). Error bars indicate standard deviation. Unadjusted *P*-values are shown; *, *P* < 0.05; **, *P* < 0.01; and ***, *P* < 0.001.

We further tested whether deletion of one *cls* gene affects the expression of the other. During logarithmic phase, loss of either gene had no detectable effect on the paralog’s transcript levels (Fig. 4B). In stationary phase, however, deletion of *clsA* caused an approximately eightfold increase in *clsB* expression, while deletion of *clsB* did not alter *clsA* levels (Fig. 4B). This asymmetric response suggests that *clsB* upregulation may partially compensate for loss of ClsA activity during stationary phase.

To determine whether membrane stress influences *cls* expression, we exposed cells to acute stress conditions for 20 min prior to RNA harvest. SDS had no significant effect on either transcript, while deoxycholate (0.01%) and elevated K+ (300 mM) each increased *clsA* expression approximately twofold and modestly decreased *clsB* expression (Fig. 4C). Acidic pH had little effect on either gene, while basic pH reduced the expression of both (Fig. 4C). Ionophore treatment produced small but statistically significant changes in both transcripts, though the biological relevance of these modest shifts during ionophore treatment remains unclear (Fig. 4C).

### Deletion of the *cls* genes does not measurably alter cellular ion homeostasis

Growth of the *cls* mutants was perturbed by elevated NaCl and KCl and by molecules that dissipate ion gradients (Fig. 2), phenotypes consistent with a compromised permeability barrier or altered ion balance. We therefore asked whether loss of *cls*-dependent lipids disrupts intracellular ion levels in *B. fragilis* P207. To address this, we used inductively coupled plasma mass spectrometry (ICP-MS), which quantifies total elemental composition and thereby provides a measure of ionic content for biologically relevant elements including Na+ and K+. Conventional cell-washing steps can artifactually perturb intracellular element levels (Fig. S5A), so we used a wash-free ICP-MS protocol adapted here for bacterial cultures (38). Under standard growth conditions in supplemented brain heart infusion (BHIS) medium, wild-type *B. fragilis* P207 cells contained approximately 550 mM phosphorus (P), 430 mM potassium (K+), 98 mM sulfur (S), and 63 mM sodium (Na+) (Fig. S5C; medium elemental concentrations are shown in Fig. S5B).

To compare wild-type and mutant strains, we normalized intracellular element levels to phosphorus content, an accepted practice in this type of analysis (39). Sulfur scaled proportionally with phosphorus across all samples (Fig. S5D and H), supporting its use as a secondary internal control. Normalizing to phosphorus revealed no significant differences in K+, Na+, or any other measurable element across the Δ*clsA*, Δ*clsB*, or double-deletion strains (Fig. 5A and B; Fig. S5C, Table S4), indicating that *cls* deletion does not overtly perturb steady-state ionic composition of the *B. fragilis* cell. We observed strain-dependent shifts in phosphorus content per CFU that were consistent with differences in OD600/CFU ratios (Fig. S5E through G); this result suggests that *cls* deletion can influence the ability of cells to consistently form colonies under standard cultivation conditions.

**Figure 5.**
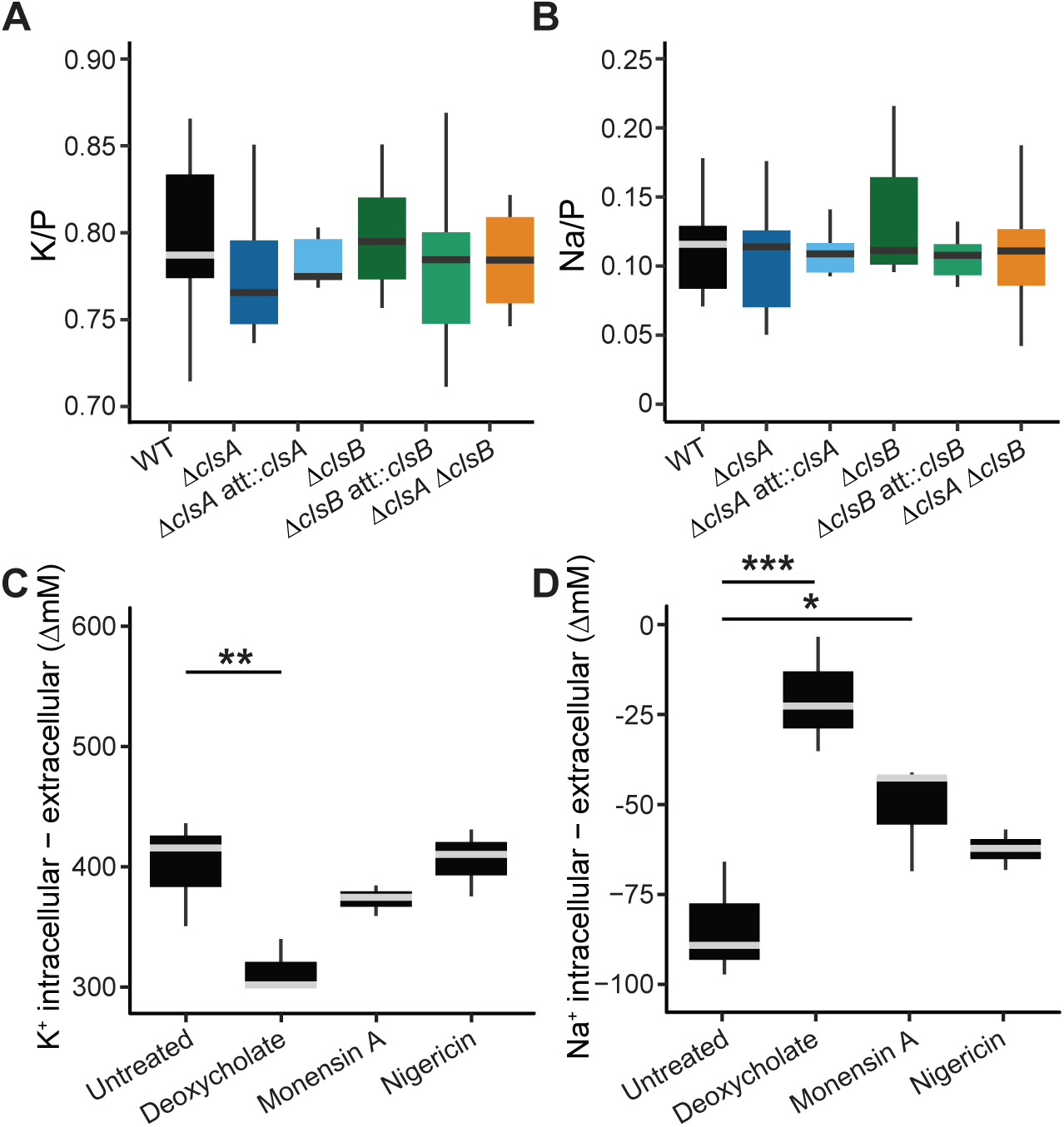
Ion homeostasis is disrupted by membrane-active stress but not by loss of *cls* function under basal conditions. Intracellular (A) potassium and (B) sodium abundance, normalized to intracellular phosphorus, in *B. fragilis* P207 WT, Δ*clsA*, Δ*clsB*, Δ*clsA*Δ*clsB*, and complemented strains as measured by inductively coupled plasma mass spectrometry. Net intracellular (C) K+ and (D) Na+ concentrations in WT cells following 20-min exposure to 0.01% (wt/vol) deoxycholate, 0.8 µg/mL monensin A, or 0.1 µg/mL nigericin. Concentrations are normalized to cell volume and colony-forming units, with extracellular concentrations subtracted; complemented strains were not included in stress treatment conditions. For all boxplots, horizontal lines indicate the median; box edges indicate the 25th and 75th percentiles; whiskers extend to 1.5× the interquartile range. *n* = 10 biological replicates for untreated conditions; *n* = 3 for stress treatment conditions. Stress-vs-untreated comparisons in panels C and D are computed against the matched-batch BHIS controls (*n* = 3). Unadjusted *P*-values are shown; *, *P* < 0.05; **, *P* < 0.01; and ***, *P* < 0.001 (linear regression).

We next asked whether membrane-disrupting treatments that impair *cls* mutant growth perturb cellular elemental content. Like most organisms, *B. fragilis* maintains a higher intracellular K+ concentration (+400 ΔmM) and a lower intracellular Na+ concentration (−67 ΔmM) than the surrounding environment (Fig. 5C and D). Exposure to 0.01% (wt/vol) deoxycholate, a microbially modified bile acid that elicits strong *cls*-dependent growth defects at this concentration (Fig. 2A) (21), caused a marked decrease in intracellular K+ and a corresponding increase in Na+ level. These reciprocal shifts in ionic composition provide direct evidence that a physiologically relevant level of deoxycholate (26) can disrupt transmembrane ion gradients in *B. fragilis* (Fig. 5C and D; Fig. S5I and J). An accompanying increase in intracellular P per CFU (Fig. S5K) suggests that some of these changes reflect the presence of non-viable cells (Fig. S5L), which retain S and P associated with large molecules such as proteins and nucleic acids while losing smaller, more diffusible ions across damaged membranes (40, 41). Consistent with its known ion selectivity, monensin A selectively increased intracellular Na+ by 32 mM without affecting K+ levels (Fig. 5C and D), mirroring the Na+ gradient disruption expected from this ionophore and aligning with the *clsB*-dependent fitness defect observed under monensin treatment (Fig. 2G). Nigericin, by contrast, produced no detectable change in K+ or Na+ under these conditions (Fig. 5C and D), suggesting either that nigericin requires longer than 20 min to act on *B. fragilis* or that cells rapidly adapt to the imposed ionic disturbance. Together, these results establish that *B. fragilis* actively maintains ionic gradients that are selectively disrupted by membrane stresses, and that *clsA/B* deletion does not alter steady-state ion composition under unstressed conditions; this is consistent with the equivalent growth of the *cls* mutants and WT in the absence of stress. The impact of ClsA and/or ClsB on ion homeostasis under stress conditions remains to be determined.

### Membrane lipid composition of WT *B. fragilis* P207

*Bacteroides* membranes contain a diverse mixture of glycerophospholipid, sphingolipid, glycosphingolipid, and peptidolipid families (23–25). To define the baseline lipid profile of *B. fragilis* P207, we used high-resolution ultra-high-performance liquid chromatography-high-resolution mass spectrometry (UHPLC-MS)-based lipidomics to characterize membrane preparations in both negative and positive electrospray ionization modes. As authentic standards were not available and because ionization efficiency varies between lipid classes, peak areas summed across different classes are not interpreted as absolute molar composition. We treat total peak area-normalized values as a semi-quantitative survey of the relative signal contributed by each lipid class. We report these values to place our findings alongside other recent Bacteroidota lipidomic studies, which also describe relative lipid-class composition (25). For making specific claims, we use OD-normalized peak areas to compare relative LC-MS signal within each ionization mode. To support more quantitative within-class comparisons, pooled samples were analyzed across a seven-point dilution series, and lipid-specific calibration curves were applied over those dilution levels (Table S3).

The WT profile contained the major lipid families reported for the *B. fragilis* group (24, 25). These include the glycerophospholipids PE, PS, and PA; the sphingolipids dihydro-ceramide and hexosylceramide; and glycine lipid, glycyl-seryl lipid, and glycyl-seryl lipids linked through a phosphate diester to diacylglycerol or dihydroceramide (Fig. 6A&6B; Fig. S6A, Table S3). The cross-class percentages reported below are semi-quantitative for the reasons given above and are intended to convey relative prominence rather than absolute abundance. Within the glycerophospholipid signal, PE species were the most prominent (39% of negative-mode peak area), followed by PS (9%), with PA and lysophospholipids contributing smaller fractions; species consistent with the mass of cardiolipins were detected as a minor class (and were dependent on the *clsB* gene) but were unsupported by MS2 (Fig. S6; Table S3). Among sphingolipid-associated signals, the most prominent were assigned to ceramide phosphoethanolamine (26% of negative-mode peak area) and dihydroceramide (7%). This is consistent with the established prominence of sphingolipids and ceramide phosphoethanolamine in *Bacteroides* membranes. Among the negative-mode WT signal from molecules whose individual chains are resolved, ∼82% is odd chain, consistent with prior work (42). Our annotations report summed acyl-chain composition and do not resolve normal-chain, iso-branched, and anteiso-branched isomers. In the *B. fragilis* group, the abundant odd-chain 15:0 and 17:0 substituents occur as both anteiso and iso forms at ratios near 1.5–2, while even-chain substituents are straight (24). Thus, a nominally saturated composition (e.g., 30:0) will likely include branched-chain species. Together, this baseline profile indicates that *B. fragilis* P207 contains the major lipid families expected for *Bacteroides* membranes and provides the reference framework for evaluating how deletion of *clsA* and *clsB* impacts cellular lipid composition.

**Figure 6.**
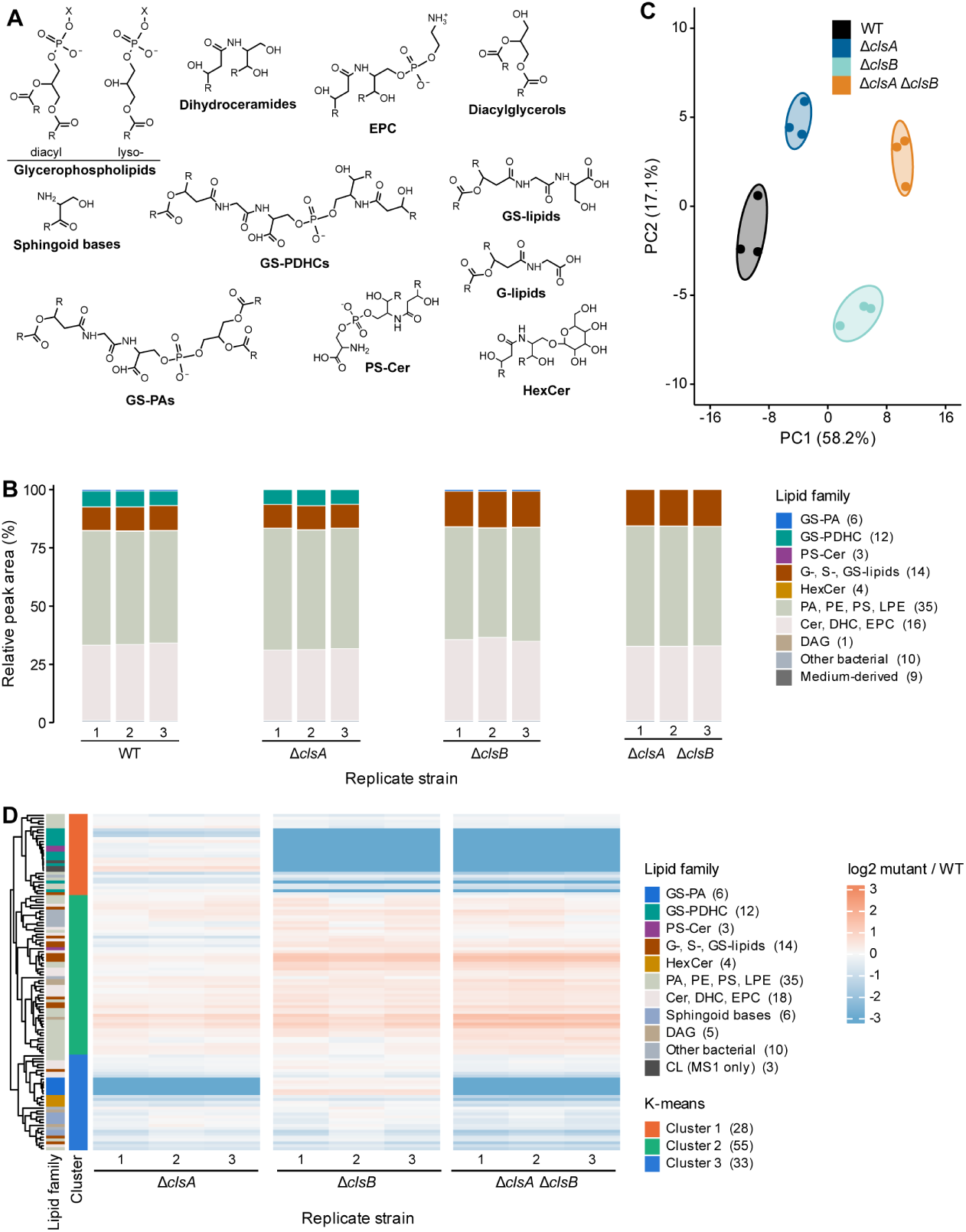
Deletion of *clsA* or *clsB* results in distinct membrane lipid changes. (A) Structures of representative lipid families detected by UHPLC-MS/HRMS with MS2 support. Family assignments are the MS-DIAL annotations, curated as described in Materials and Methods; R denotes variable hydrocarbon chains; X denotes headgroup. (B) Relative peak areas of annotated lipid families in negative ionization mode across *B. fragilis* WT, Δ*clsA*, Δ*clsB*, and Δ*clsA* Δ*clsB* strains, shown for three biological replicates, each bar summing to 100% within that mode. The number of annotated molecules in each family is given in the key. Figure S6A presents the positive-mode counterpart; peak areas are not summed across ionization modes. (C) Principal component analysis (PCA) of normalized lipid abundance profiles from WT and *cls* mutant strains. Minimum volume-enclosing ellipses were estimated using the Khachiyan algorithm. (D) K-means heatmap of relative lipid abundance (log2-transformed) in the Δ*clsA*, Δ*clsB*, and Δ*clsA* Δ*clsB* strains. To establish that these comparisons are grounded in a linear response, signal was calibrated against a pooled dilution series to compare the same lipid feature across strains (see Materials and Methods). Individual biological replicates are shown. Columns are clustered by similarity, and rows are grouped by k-means cluster. Left annotations indicate lipid family and k-means cluster. Lipid-family abbreviations correspond to the ontology fields in Table S3: Cer, ceramide; CL, cardiolipin; DAG, diacylglycerol; DHC, dihydroceramide; EPC, ceramide phosphoethanolamine; G-lipid, glycine lipid; GS-lipid, glycyl-seryl lipid; GS-PA, glycyl-seryl phosphatidic acid; GS-PDHC, glycyl-seryl phospho-dihydroceramide; HexCer, hexosylceramide; LPE, lysophosphatidylethanolamine; PA, phosphatidic acid; PE, phosphatidylethanolamine; PS, phosphatidylserine; PS-Cer, serine phosphoceramide; S-lipid, serine lipid.

### *clsA* and *clsB* mutants have distinct lipid profiles

We next compared the membrane lipid profiles of the *B. fragilis* P207 WT strain to the derived Δ*clsA*, Δ*clsB*, and Δ*clsA* Δ*clsB* mutant strains. The overall lipidome remained dominated by similar major lipid classes across strains (Fig. 6A and B), although, as noted above, individual lipid features showed genotype-dependent differences in relative signal (Fig. 6D; Table S3). We used principal component analysis as an unsupervised overview of the normalized lipid abundance data to determine whether the mutant lipidomes differed at the whole-profile level. This analysis separated WT, Δ*clsA*, Δ*clsB*, and Δ*clsA* Δ*clsB* samples, indicating that loss of each *cls* gene produces distinguishable lipidome profiles (Fig. 6C). We then used k-means clustering as an exploratory approach to identify lipid features with shared abundance patterns across the *cls* mutant backgrounds. The k-means heatmap displays individual biological replicates (Fig. 6D). The major cluster-level patterns support the conclusion that loss of *clsA* or *clsB* produces distinct lipidome changes.

Feature-level k-means clustering resolved three groups (Fig. 6D). A set of 28 molecules was strongly depleted in strains lacking *clsB*; its largest component (12 of the 28 molecules) was assigned to the GS-PDHC family (24), together with the three predicted cardiolipin-mass features. Those cardiolipin-mass features fell below detection in the Δ*clsB* and Δ*clsA* Δ*clsB* strains (Table S3). A second cluster (33 molecules) contained six molecules assigned to the GS-PA family. GS-PA species were 12- to 45-fold lower in Δ*clsA* and fell below detection in the double mutant. Both glycyl-seryl families gave family-diagnostic MS2 fragments. The remaining 55 features formed a large cluster that was retained across all mutant backgrounds and included PE, EPC, and PS. These results indicate that ClsA and ClsB make non-redundant contributions to the *B. fragilis* lipidome, with the abundance of different GS conjugate families depending on different *cls* genes.

### ClsA/ClsB and peptidolipid acyl-chain composition and the broader lipidome

The two glycyl-seryl conjugate families differ in the lipid to which the glycyl-seryl moiety is attached and in their acyl-chain distributions and apparent abundances (Fig. 7A; Table S3). The gene-lipid pattern described above is not absolute. Deletion of *clsA* reduced the GS-PDHC N-OH-acyl subfamily about twofold, consistently across all four of its molecules (*q* = 0.046), while leaving the more abundant N-O-acyl subfamily unchanged (1.03-fold; Fig. 7C). Within a subfamily, neither deletion selectively impacted particular chain lengths (Fig. S7A).

**Figure 7.**
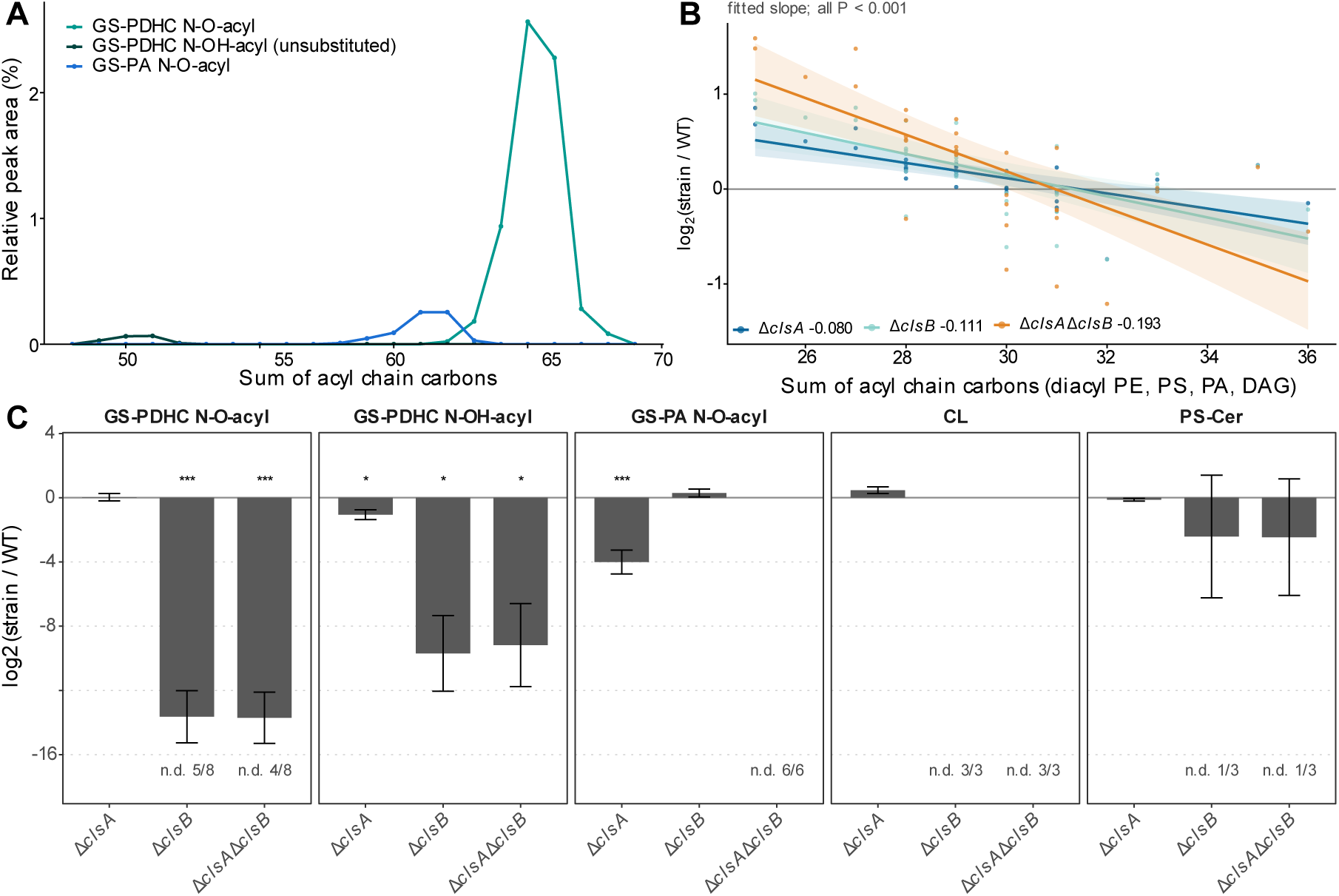
*cls* deletion produces a shared shift toward shorter acyl chains and gene-specific changes in a small number of lipid classes. (A) Chain-length distributions of the three *cls-*dependent glycylseryl classes in the wild type: GS-PDHC N-O-acyl, green; GS-PDHC N-OH-acyl, black; and GS-PA N-O-acyl, blue. Relative peak area is plotted against summed acyl-chain carbons on a common *y*-axis so that the three distributions can be compared. Negative ionization mode. (B) Log2(fold change) relative to the wild type against summed acyl-chain carbons for the diacyl glycerolipids PE, PS, PA, and DAG (25–36 carbons; 35 molecules), with ordinary least-squares fits and 95% confidence bands. Fitted slopes in log2 units per carbon are given beside each strain in the key, and each has *P* < 0.001. The double mutant differs in slope from the *clsA* single mutant (interaction *P* = 0.004); the two single mutants do not differ from one another (*P* = 0.42). (C) The five lipid classes meeting a change of at least 1.5-fold at *q* < 0.05 or falling below detection. Bars are the mean log2 ratio to the wild type over the molecules of the class detected in that genotype; error bars are the standard deviation across molecules. n.d. indicates that molecules of the class were not detected in that genotype; these are excluded from the bar rather than given a pseudocount. Where a whole class was not detected, there is no bar. Asterisks are placed directly above the bar to which they refer: *, *q* < 0.05; **, *q* < 0.01; and ***, *q* < 0.001, Benjamini-Hochberg adjusted across all classes and genotypes. All panels are from three biological replicates per genotype (Table S3). Abbreviations are as defined in the legend to Fig. 6.

Beyond the peptidolipids, the diacyl glycerolipids shifted toward shorter chains in every mutant. Regressing the log2 ratio to the wild type on summed acyl-chain length across PE, PS, PA, and DAG species of 25–36 carbons gives slopes of −0.080 in the *clsA* mutant, −0.111 in the *clsB* mutant, and −0.193 in the double mutant (each *P* < 0.001; Fig. 7B). The slope is unchanged when lipid class is included as a covariate and is negative and significant within PE, PS, and DAG separately. The two single mutants do not differ from one another (*P* = 0.42), and the double mutant differs from the *clsA* mutant (interaction *P* = 0.004) but not significantly from the *clsB* mutant. This chain-length effect is therefore shared by the two single mutants rather than specific to either and is most pronounced when both genes are absent; across the 25–36 carbon range, the double-mutant slope corresponds to about a fourfold shift in relative abundance.

One further class showed notable changes as a function of *cls* gene deletion. Serine phosphoceramide (PS-Cer) was unaffected in the *clsA* mutant, while in both strains lacking *clsB,* one of its three species fell below detection, and a second was reduced approximately 35-fold (Fig. 7C; Table S3). A *clsB* dependence therefore extends to part of a second ceramide-based class, though not to every species within it. Across the remaining lipid classes, no genotype changed the class mean by more than 2.4-fold in either direction (Fig. S7B). Together, these results indicate that ClsA and ClsB have both gene-specific and shared effects on the *B. fragilis* lipidome. The glycyl-seryl conjugates and PS-Cer depend on one gene or the other, while the shift toward shorter acyl chains occurs to a similar extent in either single mutant and is greatest when both genes are absent.

### ClsA and ClsB loss leads to distinct changes in diffusible lipids and non-lipid metabolites

We extended our analysis to ask whether the loss of *clsA* and *clsB* also differentially reshapes the broader, membrane-diffusible *B. fragilis* metabolome in addition to structural cell envelope lipids. Using LC-tandem mass spectrometry, we found that each strain had unique metabolite profiles, with *ΔclsB* being the most distinct (Fig. 8A through C; Table S5). Using publicly available databases, 56% of all reproducible features were annotated, with 83% of those matches from small molecule databases. Approximately 27% of the metabolomic features were not shared between all strains, with ∼14% of the features depleted only in *ΔclsA*, *ΔclsB*, or *ΔclsA ΔclsB* (Fig. 8B). Notably, we observed that the effect of *clsA* and *clsB* deletion on the metabolome is not additive, with *clsB* deletion showing the most profound alterations on the metabolomic profiles by hierarchical clustering (Fig. 8C).

**Figure 8.**
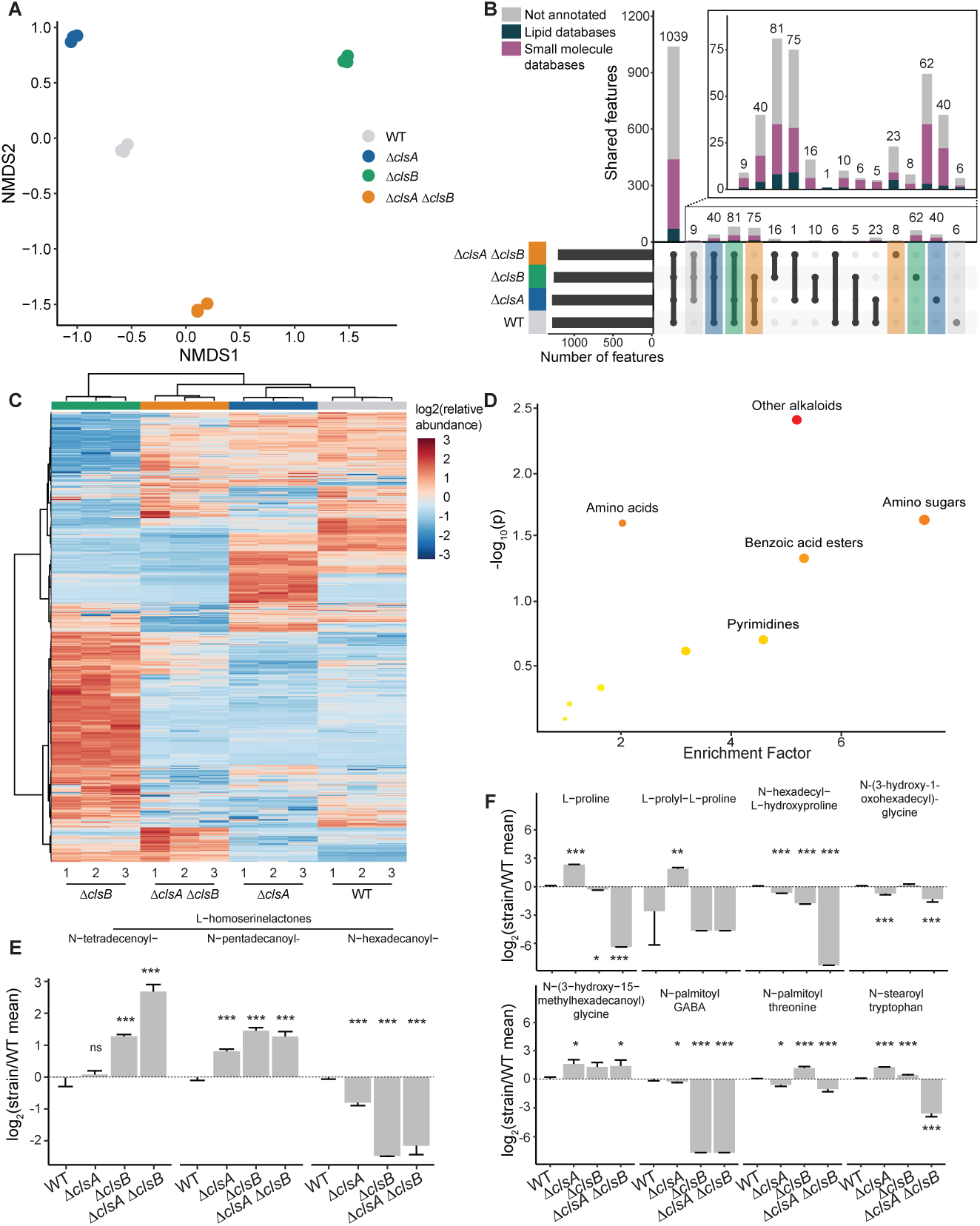
Loss of *clsA* or *clsB* function is associated with distinct metabolomic profiles. (A) Non-metric multidimensional scaling (NMDS) of metabolomics data colored by strain. (B) Upset plot with the main bar plot color-coded for the metabolomic feature annotation sources (dark green, lipid databases; purple, small molecule databases; and gray, not annotated within 5 ppm error) and the lower section color-coded for metabolomic features only present or absent in each strain (gray, WT; blue, Δ*clsA*; green, Δ*clsB*; and orange, Δ*clsA ΔclsB*). Inset is a rescaled axis of metabolites missing in at least one strain. (C) Heatmap of row-normalized abundance of each metabolite in each strain, clustering by metabolomic features on the vertical axis and samples on the horizontal axis using Euclidean distance and Ward clustering (three technical replicates are shown for each strain). (D) Metabolic pathway analysis with non-lipid sub-categories of Δ*clsAΔclsB* knockout compared to wild type (≥3 features/pathway). The *x*-axis represents the enrichment factor computed from pathway topological analysis on non-lipid species, and the *y*-axis is the log of *P*-value obtained from pathway enrichment analysis. The pathways that were most significantly changed are characterized by both a high-log(*P*) value and a high enrichment factor (top right region). (E and F) Abundance of compounds in enriched pathways for E putative *N*-acyl homoserine lactone-like features and F fatty amides (top enriched pathways) (mean and standard deviation of three technical replicates are shown; linear regression compared to WT). Across all panels, error bars indicate standard deviation. Unadjusted *P*-values are shown; *, *P* < 0.05; **, *P* < 0.01; and ***, *P* < 0.001.

To identify metabolites associated with loss of ClsA and ClsB activity, we performed pathway/chemical-class enrichment analysis comparing feature abundances between WT and the Δ*clsA* Δ*clsB* double mutant (Tables S6 and S7). Consistent with the lipidomic changes described above, fatty amides were the most differentially enriched class, followed by fatty esters and sphingoid bases (Fig. S8). In addition, non-lipid small molecule classes were also enriched, including alkaloids, amino acids, amino sugars, benzoic acid esters, and pyrimidines, several of which showed high enrichment factors and significant unadjusted *P*-values (Fig. 8D). These broad metabolic shifts indicate that ClsA and ClsB activities influence pathways beyond primary membrane architecture.

### Genotype-specific shifts in acyl amides and quorum-sensing signaling molecules

Given that amino acids and fatty amide pathways were significantly altered in the enrichment analysis, we further examined specific amino acid derivatives and diffusible lipids across the *cls* mutant backgrounds. This analysis revealed distinct, genotype-specific alterations in *N*-acyl homoserine lactone (AHL) features and *N*-acyl amides (Fig. 8E and F). AHLs and N-acyl amides are lower molecular weight, membrane-diffusible molecules with reported signaling functions (43, 44). Several of the N-acyl amides detected here are acylated amino acid conjugates, chemically related to the glycyl-seryl peptidolipids described above but not linked to a phospholipid or sphingolipid. AHLs are associated with quorum sensing (45), but gut-associated AHLs have additionally been reported to interact with mammalian intestinal epithelial and immune cells, with effects on barrier integrity and inflammatory processes (46). Mutant strains lacking *clsB* exhibited chain-length-dependent shifts in AHL-like features. Long-chain AHL-like features such as N-hexadecanoyl-HSL were depleted, while shorter-chain features such as N-tetradecenoyl-HSL and N-pentadecanoyl-HSL were enriched (Fig. 8E).

In parallel, *N*-acyl amides, a broad class of signaling lipids that can function as nutrient sources, bacterial signaling factors, or host G-protein-coupled receptor ligands (47, 48), showed distinct, non-additive alterations. While several of these features were similarly altered in both single mutants, including N-hexadecyl-L-hydroxyproline, N-(3-hydroxy-15-methylhexadecanoyl)glycine, N-palmitoyl threonine, and N-stearoyl tryptophan, others exhibited gene-specific dependency. For example, N-(3-hydroxy-1-oxohexadecyl)-glycine (i.e., commendamide) was primarily affected by *clsA* deletion, and N-palmitoyl GABA was predominantly altered by *clsB* deletion (Fig. 8F). Consistent with broader changes in this chemical space, molecular networking placed *N*-acyl amides within larger clusters of chemically related but unannotated metabolites (Fig. S9). Together, these results show that deletion of *clsA* or *clsB* influences the *B. fragilis* metabolome in ways that extend beyond membrane homeostasis, with non-additive and qualitatively distinct effects on diffusible acyl amides and AHL-like features. Given the chemical relationship between these species and the *cls*-dependent peptidolipids, these shifts may reflect redistribution of fatty acyl substrates no longer consumed during membrane lipid assembly, rather than a direct regulatory response to altered membrane composition.

## Discussion

In this study, we provide evidence that the two *B. fragilis* P207 cardiolipin synthase family proteins, ClsA and ClsB, have related but non-redundant functions in the cell. The Δ*clsA* and Δ*clsB* mutants differ in stress sensitivity, cell morphology, membrane lipid composition, and metabolite profiles, and the two genes differ in when they are regulated. The best-supported changes in the lipidome upon *clsA* and *clsB* deletion are in glycyl-seryl peptidolipid family lipids, GS-PA and GS-PDHC, respectively; both of these lipid families have MS2 support in our data set. *clsB*-dependent changes in species with accurate mass matches to select cardiolipin species are also observed, though these peaks are minor and annotation is not supported by MS2 spectra. This work thus links *clsA* and *clsB* to a class of peptidolipids that have been previously reported to influence human innate immune signaling (49, 50) and implicated in bile resistance and gut colonization in the related species *Bacteroides thetaiotaomicron* (35).

Given the known connections between membrane lipids and ion permeability, and our previous results suggesting a genetic connection between Na+/H+-dependent energy production and the *clsA* and *clsB* genes (21), we tested whether loss of *clsA* and *clsB* gene function perturbed intracellular ion homeostasis in *B. fragilis*. Under standard growth conditions, loss of *cls* function did not measurably alter intracellular K+, Na+, or other ion levels, despite the growth defects observed in *cls* mutants under osmotic and ionophore stress. The lipids that are lost as a function of *cls* deletion comprise about 8% of the negative ion mode signal, while the major components of the lipid bilayer remain intact. Our data thus provide evidence that *B. fragilis* P207 tolerates loss of GS-PDHC and GS-PA lipids (24) under standard laboratory cultivation conditions. Together, these results support a model in which ClsA and ClsB individually influence membrane composition and fitness under select stress conditions while basal ion homeostasis is maintained under standard conditions in the absence of *cls* genes (Fig. 9). Whether loss of ClsA or ClsB affects ion homeostasis under the tested stress conditions remains an open question.

**Figure 9.**
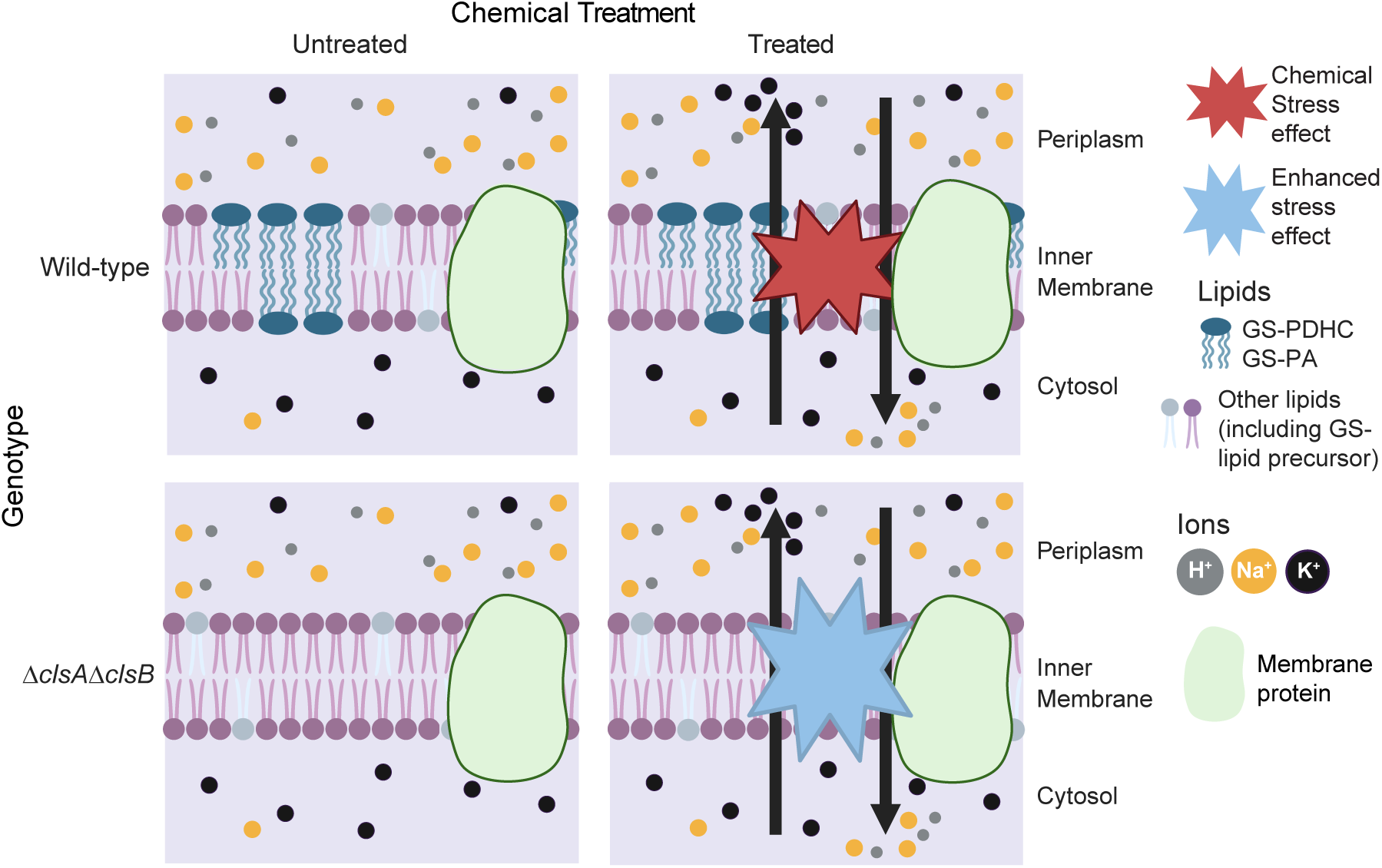
Model for ClsA/ClsB contributions to membrane composition and stress physiology. Under untreated conditions, *B. fragilis* maintains intracellular ion homeostasis, including high intracellular K+ and low intracellular Na+ relative to the surrounding medium. Loss of the *clsA* and *clsB* genes is associated with pronounced changes in membrane lipid composition; the most well-supported effects are on the glycyl-seryl lipids: GS-PA levels are strongly reduced in the Δ*clsA* mutant, and GS-PDHC is reduced to background levels in the Δ*clsB* mutant. The figure shows both GS-PA and GS-PDHC missing in the Δ*clsA*Δ*clsB* mutant; this double mutant has significant fitness loss under chemical treatments. Steady-state intracellular Na+ and K+ levels remain constant in the double mutant under standard growth conditions. Chemical perturbations, including deoxycholate and ionophores, disrupt membrane homeostasis and ion balance in wild-type *B. fragilis* P207, and these stresses have genotype-dependent effects on fitness in *cls* mutants (see Fig. 2). Together, the data support a model in which ClsA and ClsB make non-redundant contributions to membrane lipid composition and stress resilience.

Metabolomic changes in *cls* mutants extend beyond the membrane lipidome and differ between Δ*clsA* and Δ*clsB*. The genotype-specific alterations in *N*-acyl amides and AHL-like features are particularly notable, as these compound classes include molecules implicated in bacterial signaling and host interaction (46–48, 51). Some of these changes may reflect redistribution of fatty acyl substrates that are no longer channeled into membrane lipid biosynthesis, rather than direct regulatory consequences of altered membrane composition. Regardless of mechanism, this broader pattern indicates that ClsA and ClsB are connected (directly or indirectly) to aspects of *B. fragilis* metabolism that extend beyond membrane lipid homeostasis and have the potential to impact mammalian physiology.

The substrates and products of ClsA and ClsB remain to be defined biochemically, but the lipidomic data point to a testable model. GS-PA and GS-PDHC carry the same glycyl-seryl headgroup as the free GS-lipids, which accumulate in backgrounds lacking *clsB*; in both conjugates, that headgroup is joined to its lipid through a phosphate. One possibility is that these phospholipase D superfamily proteins use the free serine hydroxyl of the GS-lipid as the acceptor for lipid transfer, with ClsA transferring a phosphatidyl group and ClsB a phosphoryl dihydroceramide group. Distinguishing this from alternative models will require purified enzymes and defined substrates. Whether the lipids that depend on these genes impact fitness in the mammalian gut environment is also an open question. In *B. thetaiotaomicron*, loss of the glycine lipid synthase *glsB* removes both glycine and glycylseryl lipids, strongly raises bile sensitivity, and impairs colonization of germ-free mice (35). It remains to be tested whether the lipids that depend on *clsA* and *clsB* contribute to gut fitness in the same way. The mutant strains and lipidomic data set reported here provide a starting point for this future work.

## Materials And Methods

### Bacterial growth and strain construction

#### Strain generation

*E. coli* strains were grown aerobically at 37°C in Miller LB broth. As appropriate, *E. coli* cultures were supplemented with filter-sterilized 300 µM diaminopimelic acid (Sigma-Aldrich) to grow auxotrophic strains and/or 100 μg/mL carbenicillin to select for plasmid-containing cells. All *B. fragilis* strains were grown in supplemented brain heart infusion medium. As appropriate, cultures were supplemented with 5 μg/mL erythromycin for vector selection, 100 ng/mL anhydrotetracycline (aTC) for inducible ssbfe1 counter-selection (52), and 50 ng/mL aTC to induce expression from aTC-inducible complementation vectors (52). *B. fragilis* strains were manipulated aerobically and grown at 37°C in an anaerobic chamber (Coy Laboratory Products) filled with 5% CO2, 2.5% hydrogen, and 92.5% nitrogen.

Mutant strains were generated by allelic exchange using modified vectors described previously (52). For conjugation into *B. fragilis* P207, plasmids were transformed into the donor *E. coli* strain WM3064. Allele exchange was conducted as previously described (52). The Δ*clsA* mutant (lacking ptos_000612) was complemented by integration of an aTC-inducible *clsA* construct. The Δ*clsB* mutant (lacking ptos_003252) was complemented by integration of *clsB* under the control of a constitutive *Bacteroides* promoter (P*rpoD*, bt_1311*/*BT_RS06635). Complementation constructs were integrated into the chromosome at the *att* site using an intN1 integrase (53). Primers, vectors, and strains are listed in Table S1. See supplemental methods for additional details.

#### Cls conservation analysis

##### Alignment

The *B. fragilis* P207 ClsA and ClsB gene and protein sequences, as well as *E. coli* K-12 MG1655 (NC_000913) ClsA, ClsB, and ClsC, were aligned using default settings of the Geneious alignment algorithm (Geneious Prime 2024).

##### DeepTMHMM

The ClsA and ClsB amino acid sequences from *B. fragilis* P207 and *E. coli* K-12 MG1655 were uploaded to DeepTMHMM (v. 1.0.42, https://dtu.biolib.com/Deep TMHMM) to predict transmembrane domains (54).

##### Inter-phylum tree

Representative taxa were selected from the top five phyla in the human gut (metadata listed in Table S2) (55). Annotated *cls* genes were obtained from genomes on NCBI, and proteins were aligned in Geneious. One genome per species was used, aside from the two *B. fragilis* strains. Trees were reconstructed by neighbor joining in Geneious Prime 2024 from the protein alignments described above, using the Jukes-Cantor distance correction. Branch support was assessed by bootstrap resampling with 100 replicates and a 50% consensus support threshold; support values are shown in Fig. 1 where they exceed 80% and in Fig. S1 where they exceed 90%.

##### Intra-Bacteroides tree

TIGR04265 domains in *B. fragilis* ClsA and ClsB were identified using the conserved domain database of NCBI (56). *Bacteroides* protein sequences containing this domain were obtained by querying the InterPro database. Alignment, tree reconstruction, and tree visualization were performed as with the inter-phylum tree, except that *E. coli* K-12 MG1655 ClsA was specified as an outgroup.

##### Genomic neighborhoods

WebFlags2 was used to query genomic neighborhoods around *B. fragilis* P207 *clsA* and *clsB* genes (57).

### Growth curve assays

Stationary-phase *B. fragilis* cultures were back-diluted to an OD600 of 0.05 and grown to an OD600 of 0.3 in BHIS containing 50 ng/mL anhydrotetracycline. For growth-curve assays that included aTc-inducible complementation strains, all strains being compared, including WT, deletion mutants, and complemented strains, were grown with the same concentration of aTc to control for any effect of the inducer. Cultures were then diluted to an OD600 of 0.025–0.05 in 96-well plates containing the indicated stress condition and 50 ng/mL aTc. OD600 was measured with a Tecan Infinite M Nano plate reader for 24 h. Three biological replicates, each in technical triplicate, were performed for each growth condition.

To expose cells to stress conditions, BHIS medium was prepared as described above and then modified for each assay. For pH assays, fully supplemented BHIS was adjusted to the indicated starting pH. No additional buffering agent was added; therefore, this assay tested growth after exposure to medium adjusted to an initial pH rather than growth under actively maintained constant-pH conditions. For deoxycholate and SDS assays, 0.01% (wt/vol) deoxycholate (Sigma) or 0.002% (wt/vol) SDS (Lab Scientific) was added to cultures from filter-sterilized 1% (wt/vol) stocks dissolved in ultrapure water. For Na+ and K+ assays, NaCl (Fisher Scientific) and KCl (Fisher Scientific) were dissolved to a 4× concentration of each cation in BHIS medium and then filter sterilized before being added to cells to yield the final concentration of cells and cation (i.e., 300 mM). To determine how much additional NaCl or KCl to add, the average Na+ and K+ content of our BHIS medium was determined using ICP-MS (see below for methods), yielding 130 mM for Na+ and 33 mM for K+ (Table S4). For ionophore assays, a concentrated ionophore (monensin A or nigericin; Sigma-Aldrich) was added to cultures to yield final concentrations of 0.8 µg/mL monensin A or 0.1 µg/mL nigericin, with equivalent volumes of ethanol added to non-treated conditions as a vehicle control. We calculated maximum OD600 and doubling time using R (ipolygrowth; see supplemental Methods) (58).

### Microscopy

*B. fragilis* P207 cell cultures were grown overnight under antibiotic selection and then back-diluted into medium containing inducer (aTC) without antibiotic selection to reach log-phase growth. All cultures were removed from the 37°C incubator but were maintained under anaerobic conditions, with only two or three cultures removed for microscopy at a time. Cells were visualized by phase-contrast microscopy on a Leica DMI6000 B inverted microscope with a Hamamatsu ORCA-R2 10600 camera with a 63× Plan Apo objective. Each strain was visualized on four different days to generate four biological replicates. The two images per replicate from each strain/day were chosen for analysis. Chosen images were processed using the MicrobeJ plugin (v. 5.13I) cell size analysis software in Fiji (v. 1.54k) (59). Called cells were manually curated, and resulting cell lengths and widths were exported and analyzed in R to calculate cell volume and Epanechnikov kernel densities.

### Gene expression assays

#### Stress condition exposure

For comparison of *cls* gene expression at log and stationary phases, *B. fragilis* WT, *ΔclsA*, and *ΔclsB* strain log-phase cultures were back diluted to an OD600 of 0.05 without inducer. At 2 and 24 h, cell samples were pelleted and resuspended in 1 mL of TRIzol (Invitrogen). For acute toxicity assays, back-diluted WT *B. fragilis* cultures that reached OD600 ∼0.3–0.5 were exposed to a specified condition (i.e., pH, SDS, deoxycholate, K+, monensin A, or nigericin, as described for the growth assays) and incubated for 20 min before cell sample collection.

#### RNA extraction

RNA was extracted as described previously (21). RNeasy Extraction Kit (Qiagen) was used to purify samples treated with Turbo DNase (Invitrogen) using the manufacturer’s protocol.

#### RT-qPCR

Samples were run in technical triplicate. Ct values were determined using a QuantStudio5 (Applied Biosystems) instrument and a Luna One-Step RT-qPCR Kit (NEB) for target genes in each sample (21), using *B. fragilis dnaN* (ptos_002600, encoding DNA polymerase subunit beta) as a normalization gene.

### Inductively coupled plasma mass spectrometry

Log-phase cells were removed from the anaerobic chamber for sample collection. If exposed to a stress, cells were incubated for 20 min aerobically at room temperature with either 0.01% (wt/vol) deoxycholate, 0.8 µg/mL monensin A, or 0.1 µg/mL nigericin. Cultures were spiked with gadolinium-DOTA (Gd-DOTA, Macrocyclics) to a final concentration of 40 µM. Washing cells, even with the BHIS medium in which the cells were grown, perturbs intracellular concentrations of small alkali metals such as Na+ and K+, making quantification difficult (Fig. S5A). Using a Gd-DOTA spike enabled us to remove washing steps in the ICP-MS sample preparation process (60, 61).

For the wash experiment, wash condition cells were washed twice with either 250 mM sucrose or BHIS medium. Then, unwashed cells (i.e., no previous centrifugation) and resuspended washed cells were transferred to a metal-free tube, given a Gd-DOTA spike to 40 µM, and then pelleted. For *cls* and treatment comparison experiments, cells were unwashed.

Weighed cell pellets and supernatant were dried overnight and then digested using 70% HNO3 acid, before diluting to 3% nitric acid. All standards, blanks, and ICP-MS samples were prepared using ultra-trace metal grade nitric acid (70%, Fisher), ultrapure water (Millipore), metal-free polypropylene conical tubes (15 and 50 mL, Labcon), and trace metal grade pipette tips (Labcon). All solutions were weighed for accurate elemental determination using the XSR205 DU semi-analytical balance (Mettler Toledo).

Samples were analyzed using an Agilent 8900 Triple Quadrupole ICP-MS (Agilent Technologies). To prepare ICP-MS standards, the multi-element standard IV-65024 (Inorganic Ventures) was diluted with 3% (vol/vol) nitric acid in ultrapure water. Internal standardization was accomplished inline using a 200 ppb internal standard solution in 3% (vol/vol) nitric acid in ultrapure water (IV-ICPMS-71D, Inorganic Ventures). The isotopes selected for analysis were 23Na, 31P, 32S, 39K, and 157Gd with 45Sc, 89Y, 115In, and 159Tb used for internal standardization. For accurate quantification of 31P and 32S, these isotopes were measured in the oxygen mode. To quantify intracellular elemental concentration, the average strain-specific cell volume was determined and then the molar concentration calculated for each element from the number of atoms per cell. The ΔmM concentrations were calculated by subtracting the concentration of an element in the medium from that in the intracellular space.

### Lipidomics assays

WT, Δ*clsA*, Δ*clsB*, and Δ*clsA* Δ*clsB* strains were backdiluted, grown to OD600 of 0.7, and then submitted on dry ice to the University of Tennessee-Knoxville Biological and Small Molecule Mass Spectrometry Core (RRID: SCR_021368) for targeted lipidomic analysis of membrane lipids. Lipids were extracted as previously described (see supplemental Methods for details) (62). The lipidomic analyses were performed using a previously validated method (62) on an ultra-high-performance liquid chromatography system coupled to a high-resolution mass spectrometer. The chromatographic separations were carried out using a Vanquish Horizon LC system (Thermo Scientific) and reversed-phase separations as described previously (62). The eluent was introduced to an Exploris 120 mass spectrometer (Thermo Scientific) via electrospray ionization for high-resolution analysis. Following acquisition, lipid features were annotated using MS-DIAL (31). The peak areas reported in Table S3 were obtained by re-integrating the raw files with a peak-detection, alignment, and adduct-grouping workflow developed for this study, in which each molecule is quantified as the sum of its ion forms within a single ionization polarity rather than from a single adduct. MS-DIAL annotation was performed separately for positive- and negative-ion modes using an MS1 tolerance of 0.01 Da, an MS/MS tolerance of 0.025 Da, and a 2-min retention-time tolerance. Candidate features were filtered against blank samples, and features with incorrect exact mass assignments or ghost peaks were removed, including by using the MS-DIAL/MS-CleanR workflow after initial annotation. Because many detected lipid features lacked MS/MS spectra, annotations were manually reviewed; retained features were required to be within 5 ppm of the expected exact mass, and MS/MS spectral matches were manually inspected when available. Peaks were retained only when the reported signal-to-noise ratio was ≥3. Thus, lipid assignments should be considered annotations based on LC-MS, supported by accurate mass, retention behavior, blank filtering, and manual review, with MS/MS support where available.

To account for feature-specific differences in MS response, pooled lipid extracts were analyzed across a seven-point serial dilution series spanning six orders of magnitude (1×– 106× dilution, in triplicate). Most lipid features were detected only at the two or three least-diluted levels (1×, 10×, and 100×); features were retained for quantification when they were detected at a minimum of two consecutive dilution levels in at least two of three replicates per level. For each retained feature, a calibration curve was fit by linear regression of integrated peak area against relative lipid amount, where relative amount was defined as the reciprocal of the dilution factor (1/dilution). For each experimental sample, a calibration-curve-derived “amount equivalent” was then calculated for each lipid feature as (area − intercept)/slope, expressed in units of the original pool concentration. Sample amount equivalents were further normalized to the mean amount equivalent in WT samples to express each lipid in *cls* deletion strains as a fold change relative to WT. Because authentic standards were not available for all annotated lipid classes, these normalized values were treated as semi-quantitative rather than absolute measurements of lipid abundance.

### Metabolomic assays

WT, *ΔclsA*, *ΔclsB*, and *ΔclsA ΔclsB* strains were grown in 50 mL BHIS medium, and metabolites were extracted with ethyl acetate (see supplemental Methods for details). Untargeted metabolomics data were acquired using a Bruker timsTOF Pro2 (Bruker-Daltonics, Billerica, MA, USA) coupled to an Agilent 1290 Infinity II Bio UHPLC (Agilent, Santa Clara, CA, USA). The chromatographic separations were carried out using an Acquity UPLC HSST3 column (2.1 × 150 mm, 1.8 μM) (see supplemental Methods for details). Following data acquisition, mass spectrometry data were pre-processed using Bruker MetaboScape version 9.0.1 (Bruker-Daltonics, Billerica, MA, USA) and filtered using mpactR (see supplemental Methods for details) (63). Peak intensity for each metabolite was normalized to the total peak intensity of all features detected in the sample. Molecular networking was generated as previously described (64) using Global Natural Products Social Molecular Networking 2 (http://gnps2.org/). A minimum cosine similarity score of 0.7 with at least six matching peaks, a parent mass tolerance of 2 Da, and a fragment ion tolerance of 0.5 Da were selected to generate consensus spectra. Files were imported into Cytoscape 3.10.3, and nodes were arranged with the yFiles organic layout plugin (65). Pathway analysis was performed with MetaboAnalyst 6.0 functional analysis [LC-MS] module (66). A formatted peak intensity table was uploaded to the web server (https://metaboanalyst.ca) and then normalized by sum and log transformed. The default setting of 5.0 ppm mass tolerance and the mummichog algorithm v2.0 with a *P*-value cutoff of 0.0005 were used against the lipid and non-lipid pathway libraries, with at least three entries in each pathway. Metabolites from enriched pathways were plotted using R (v4.5.0).

### Data analysis and reproducibility

All data analyses were performed using R (v4.3.3 unless otherwise specified) and several packages (58, 67–91). Code used and original data sets for all data analysis and figure generation, except for the WebFlags neighborhoods, are published on GitHub (https://github.com/mschnizlein/bfrag_cardiolipin).

## Acknowledgments

We thank the Biological and Small Molecule Mass Spectrometry Core (BSMMSC) at the University of Tennessee-Knoxville and the Q-Beam Center at Michigan State University for their work processing and analyzing samples for lipid and elemental analyses, respectively. We thank Drs. Kiwon Ok, Keith MacRenaris, and Aaron Sue from the O’Halloran Lab for their advice on using the ICP-MS and analyzing output. We thank the entire Crosson Lab for their advice as the project progressed, particularly Dr. Melene Alakavuklar for her feedback on this manuscript and figure presentation. Dr. Yolanda Huang provided helpful questions and comments on a preprint of this manuscript that spurred additional analysis, and the anonymous reviewers provided thoughtful questions and critical feedback that greatly improved the manuscript. We used BioRender to generate the model presented in Fig. 9. We thank two undergraduates who piloted several experiments: Damone Charleston and Timothy Baldwin. We thank Kathryn McBride for strain culturing in the Balunas Lab. *E. coli* strain WM3064 was a gift from William Metcalf at UIUC. Work on this project in the S.C. lab was supported in part by NIDDK (RC2DK122394) and NIGMS (R35GM131762), in the T.V.O. lab by NIGMS (P41GM135018 and R01GM038784), and in the M.J.B. lab by NCI (U01CA264071).

## Data Availability

All data supporting the findings of this study are available within the article and its supplemental material. Lipidomics data and associated metadata were deposited in MetaboLights under accession MTBLS11891 (92). Untargeted metabolomics data were deposited in the Mass Spectrometry Interactive Virtual Environment (MassIVE) under accession MSV000102762. Processed ICP-MS, lipidomics, and metabolomics data sets are additionally provided in Tables S3 to S7.

