## supplemental figures & methods for "Non-redundant cardiolipin synthase family proteins shape lipid composition and stress resilience in *Bacteroides fragilis*"

### Supplemental Figures and Legends

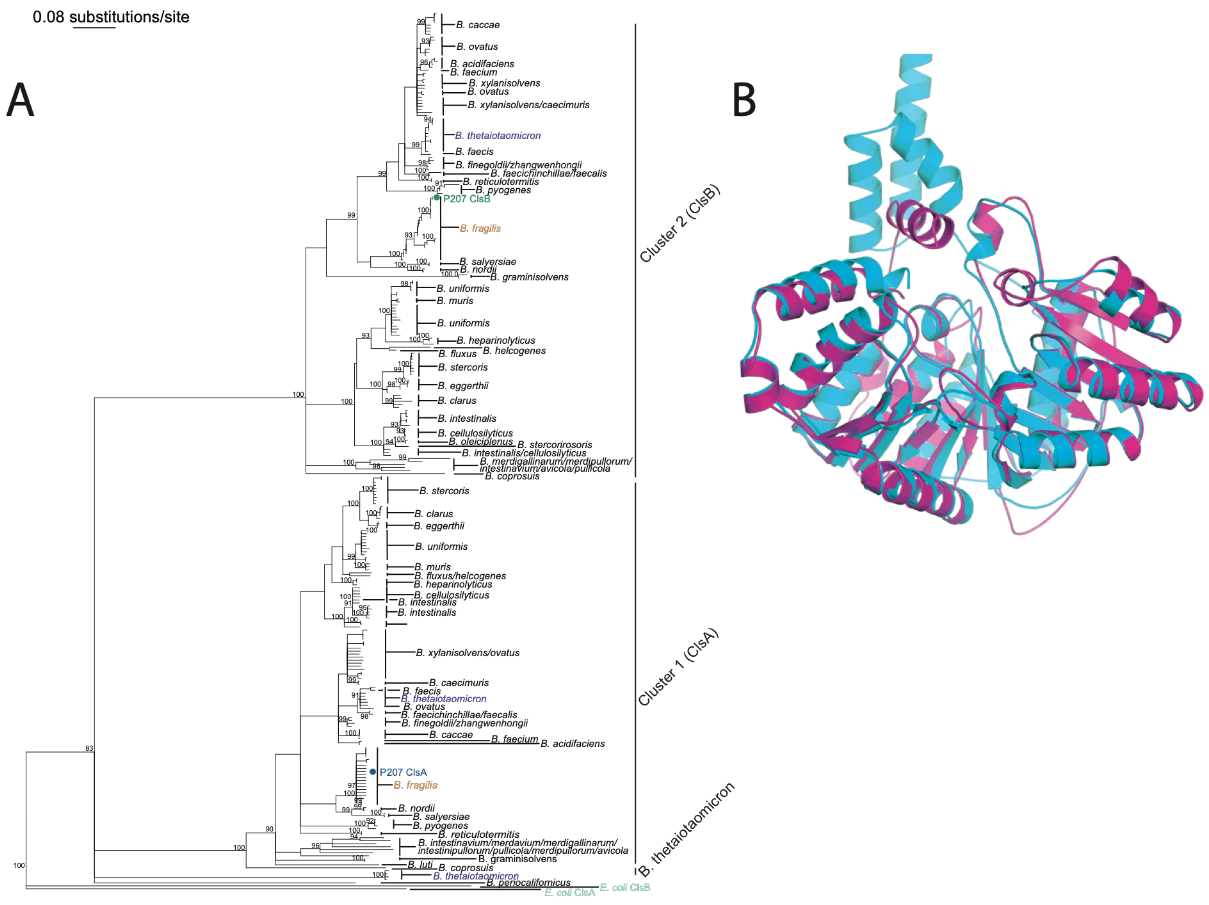


**Figure S1. Neighbor-joining tree of cardiolipin synthases from *Bacteroides* species. (A)** Clusters 1 (ClsA), 2 (ClsB) and 3 (not named) of *Bacteroides* Cls proteins are indicated. *E. coli* ClsA was designated as an outgroup. ClsA and ClsB of *B. fragilis* P207 are colored in blue and green, respectively. Other strains of *B. fragilis, B. thetaiotaomicron*, and *E. coli* are colored in gold, purple and light blue, respectively. Taxa labels with similar tree positions are combined for clarity at times (i.e., *B. xylanisolvens/caecimuris*). Key indicates the rate of amino acid substitutions per site. Bootstrap values are generally shown only if ≥90; as in Fig. 1 support is based on 100 bootstrap replicates. **(B)** Structural overlay of Alphafold3 predicted structures of ClsA and ClsB of *B. fragilis* P207; ClsB (pTM=0.91), ClsA (pTM=0.89). ClsA in cyan, ClsB in magenta. Structural fit RMSD = 0.93 Å across 320 residues. The N-terminal hydrophobic region of ClsA (visible at the top of the panel) adopts a structurally distinct conformation from the corresponding ClsB region.

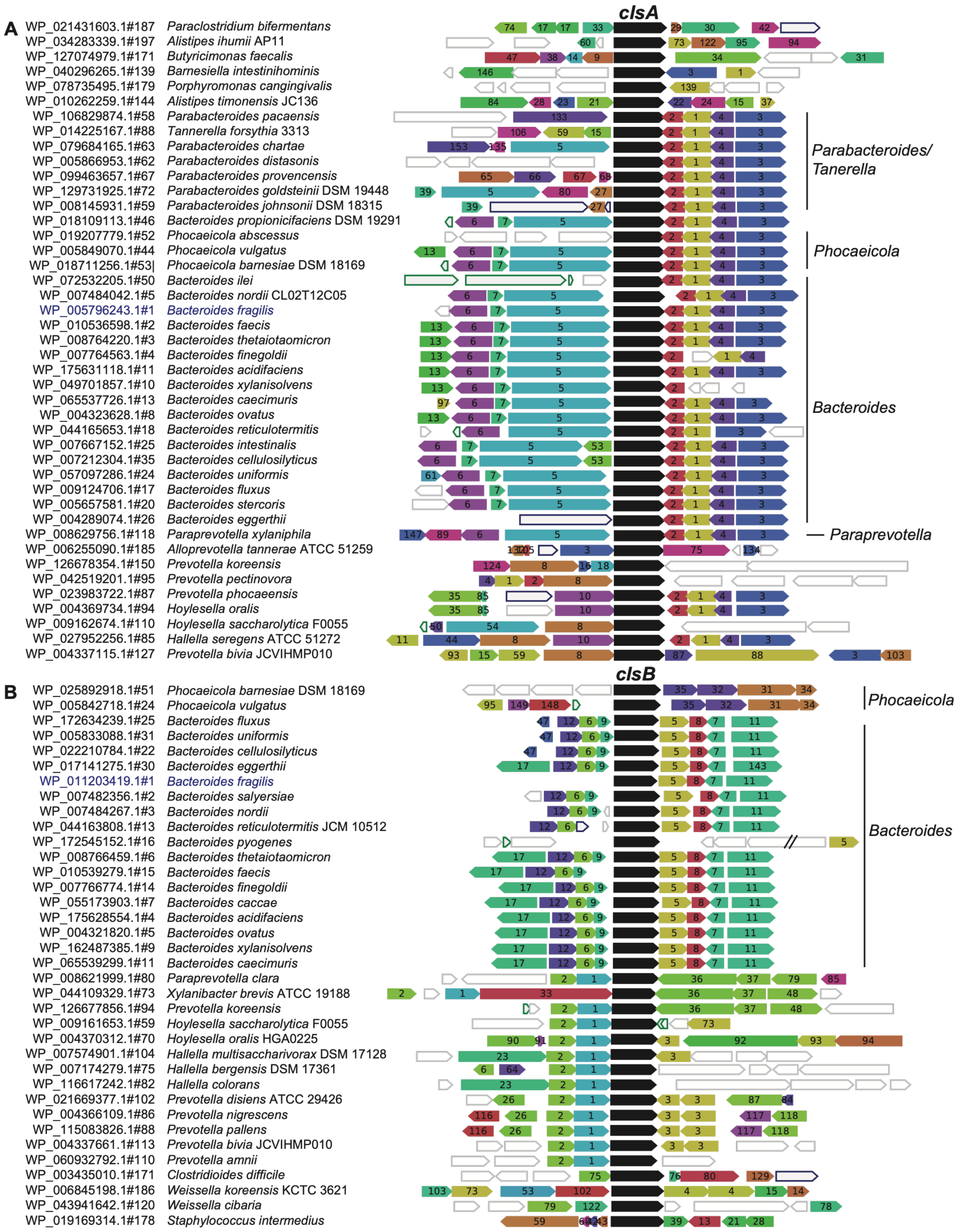


**Figure S2. Conserved genomic neighborhoods distinguish *Bacteroides clsA* and *clsB* loci.** Curated WebFlags2 output of (**A.**) *clsA* and (**B.**) *clsB* genomic neighborhoods. *B. fragilis* is colored blue. Genera of interest with similar levels of gene conservation are indicated at right.

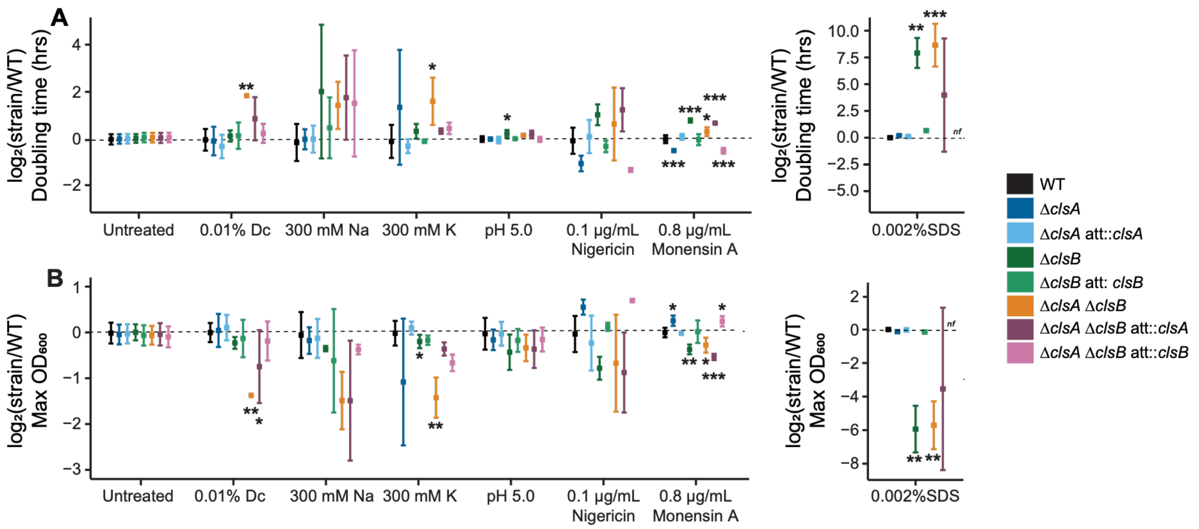


**Figure S3. Loss of cardiolipin synthase family genes results in growth defects under stress. A., B.** $\log_2$(fold change) of (**A.**) doubling time and (**B.**) maximum $OD_{600}$ comparing *B. fragilis* wild-type (WT) and *cls* mutants to the average of WT in that condition. Metrics were calculated from 24-h growth curves in BHIS medium alone or containing 0.01% deoxycholate (Dc), 0.002% sodium dodecyl sulfate (SDS), 300 mM sodium ($Na^+$), 300 mM potassium ($K^+$), and 0.1 µg/mL nigericin and 0.8 µg/mL monensin A. Any doubling time ≥100 h was considered as no growth. The dashed horizontal line indicates no change from the average of WT. Strain are colored with blue (*clsA-related*), green (*clsB*-related) and orange/purple (*clsA clsB*-related). Black indicates WT. The vertical lines indicate mean; the ends of the whiskers ± standard deviation (linear regression compared to WT). Three biological replicates for each condition were run, each run in technical triplicate. nf, no accurate fit of growth data by the polynomial was found. Unadjusted p-values are shown; *, $p < 0.05$; **, $p < 0.01$; ***, $p < 0.001$.

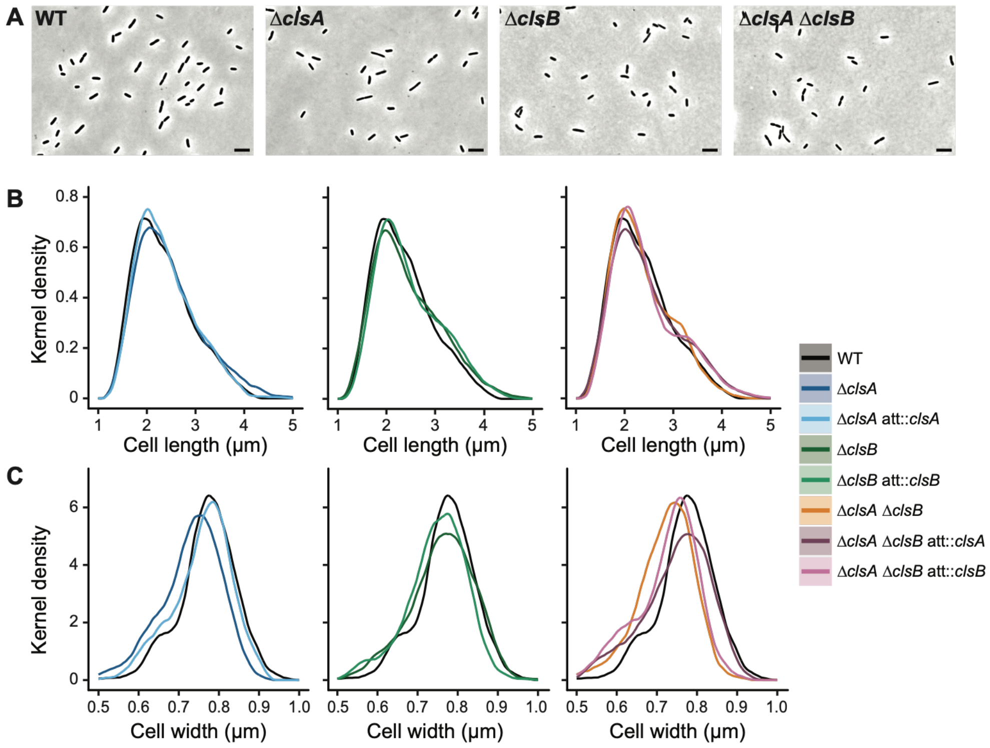


**Figure S4. Loss of *cls* genes is associated with changes in cell shape. A.** Phase-contrast microscopy images of *B. fragilis* P207 WT, Δ*clsA*, Δ*clsB* and Δ*clsA* Δ*clsB* strains. Scale bars, 5 μm. **B., C.** Distributions estimated by kernel density of (**B.**) cell length and (**C.**) cell width. Results are plotted by strain with blue, green and orange/purple indicating *clsA*-, *clsB*- and *clsA clsB*-related strains, respectively. Black indicates WT.

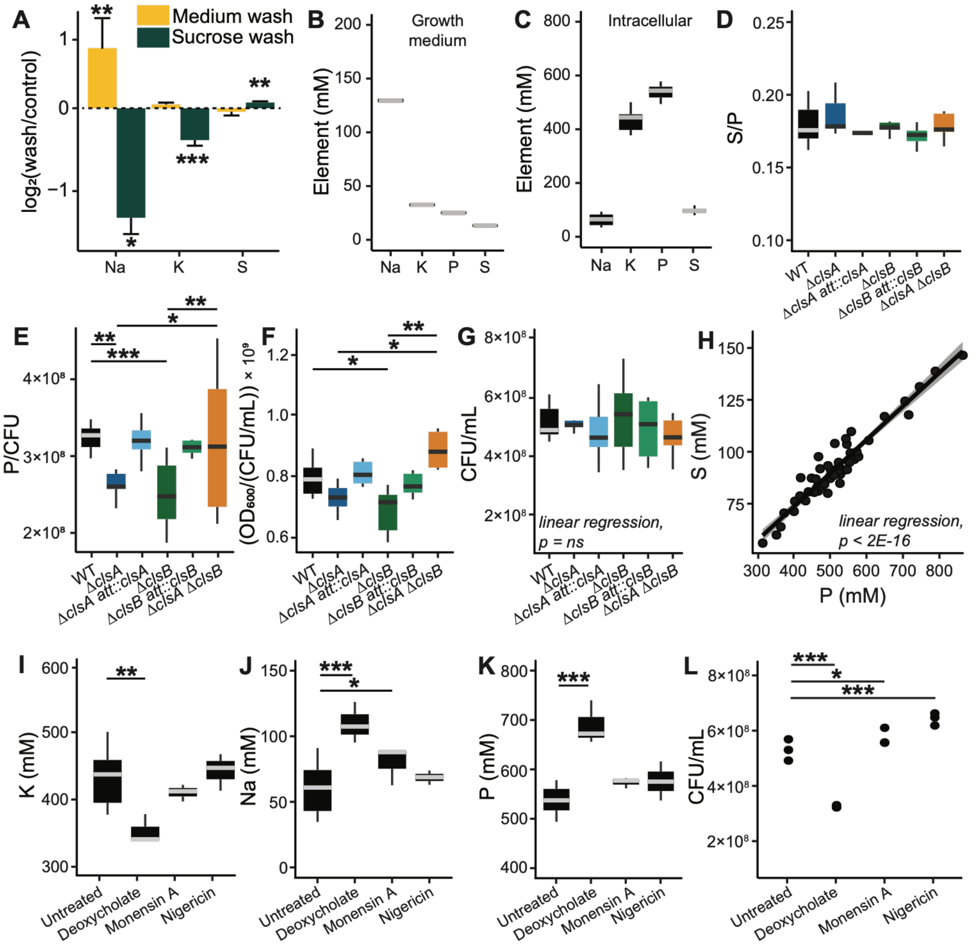


**Figure S5. Washing but not loss of *cls* genes disrupts intracellular elemental concentrations.** Inductively coupled plasma mass spectrometry (ICP-MS) analysis using wash-free sample preparation to quantify intracellular elemental concentrations. **A.** $\log_2$(fold change) of elements comparing unwashed cells to cell pellets washed with either BHIS growth medium or an isotonic sucrose solution. Error bars indicate standard deviation. **B.** Concentrations of detected elements in BHIS growth medium. **C.** Intracellular concentrations of detected elements in WT *B. fragilis* P207. **D.** Intracellular sulfur in *B. fragilis* P207 strains normalized to P. **E.** Intracellular phosphorus (P) per $10^6$ colony-forming units (CFU) in *B. fragilis* P207 wild-type (WT), Δ*clsA*, Δ*clsB* and Δ*clsA* Δ*clsB* strains. **F.** $OD_{600}$ normalized to CFU per mL of strain cultures used in ICP-MS input. **G.** Raw CFU/mL of each strain shown in panel F. **H.** Intracellular P content compared to sulfur (S) content across *cls* strains and treatment conditions. **I., J.** Intracellular concentrations of (**I.**) $K^+$ and (**J.**) $Na^+$ after a 20 min exposure to 0.01% deoxycholate, 0.8 µg/mL monensin A and 0.1 µg/mL nigericin. Each treatment had the same culture input. **K.** Intracellular P concentrations of *B. fragilis* WT cells after 20 min of exposure to the indicated stress conditions. **L.** CFU/mL of strain cultures after exposure to the treatments in **I–K**. For all boxplots, the gray or black middle lines indicate the median; the top and bottom edges of the box indicate the $25^{th}$ and $75^{th}$ percentiles of n = 3 replicates for wash and treatment experiments and n = 10 replicates for rest. Whiskers indicate 1.5× the interquartile range (IQR). For the linear regression, the shade ribbon indicates standard error of the mean. All statistical comparisons were made using linear regression, with unadjusted p-values shown; *, $p < 0.05$; **, $p < 0.01$; ***, $p < 0.001$.

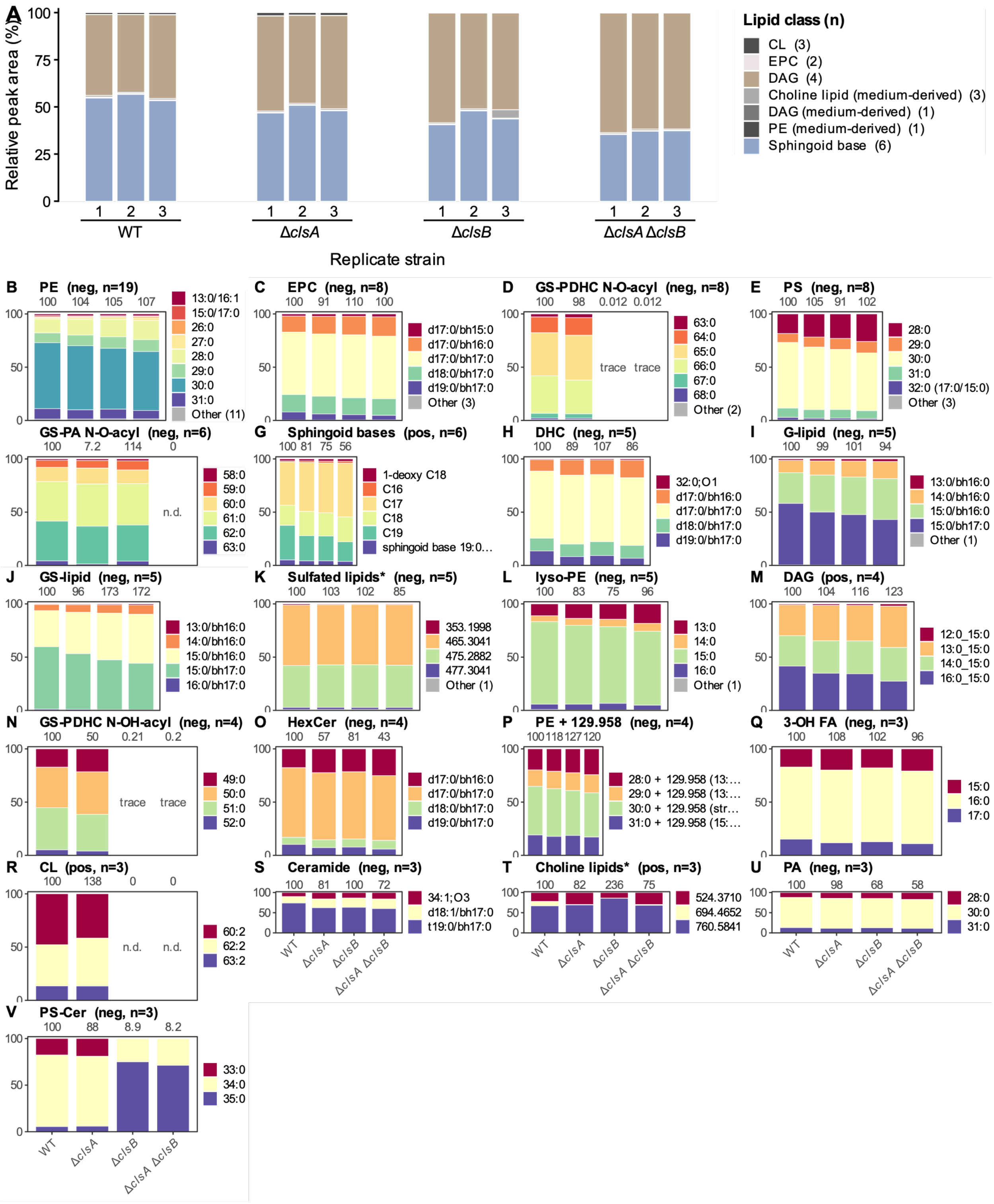


**Figure S6. The impact of *B. fragilis* P207 cardiolipin synthase family (*cls*) genes on membrane lipids.** Relative peak area of annotated lipid species in membranes across *B. fragilis* WT, Δ*clsA*, Δ*clsB*, and Δ*clsA* Δ*clsB* strains. Positive- and negative-mode data are kept separate throughout and are never summed, as relative response differs between ionization modes; **A** is positive mode, and the ionization mode of each of B-V is given in its panel title (pos, positive; neg, negative). **A.** Relative peak area of each annotated lipid class in positive ionization mode, shown for three biological replicates, each bar summing to 100% within that mode. Colors designate lipid class, and the number of annotated molecules in each class is given in the key; Fig. 6B is the negative-mode counterpart of this panel. B-V. Species composition within each lipid class, shown as the mean of three biological replicates per strain. Every class with at least three annotated molecules in one ionization mode is shown, and the 21 panels together account for 114 of the 130 annotated molecules. Species below 1% peak area in every strain are colored gray. As each bar is normalized to 100%, the bars show composition only; the figures above each panel give that strain's class total as a percentage of WT, so that a change in the abundance of a class is visible alongside any change in its composition. **n.d.,** class not detected in that strain; **trace**, class present at 2% or less of the level in the genotype in which that class is highest. Asterisks denote medium-derived classes. Strain labels are shown once at the foot of each column and apply to all panels above them (same order left to right). Lipid class abbreviations: PA, phosphatidic acid; PE, phosphatidylethanolamine; lyso-PE, monoacyl PE; PS, phosphatidylserine; CL, cardiolipin; DAG, diacylglycerol; Cer, ceramide; DHC, dihydroceramide; CerP, ceramide phosphate; EPC, ceramide phosphoethanolamine; HexCer, hexosylceramide; PS-Cer, serine phosphoceramide; G-lipid, S-lipid and GS-lipid, glycine-, serine- and glycyl-seryl-conjugated lipids; GS-PA, glycyl-seryl phosphatidic acid; GS-PDHC, glycyl-seryl phospho-dihydroceramide; 3-OH FA, 3-hydroxy fatty acid. N-O-acyl and N-OH-acyl denote the two amide-linked GS-PDHC and GS-PA subclasses. Molecule identities, ionization modes and quantitative values are in Table S3.

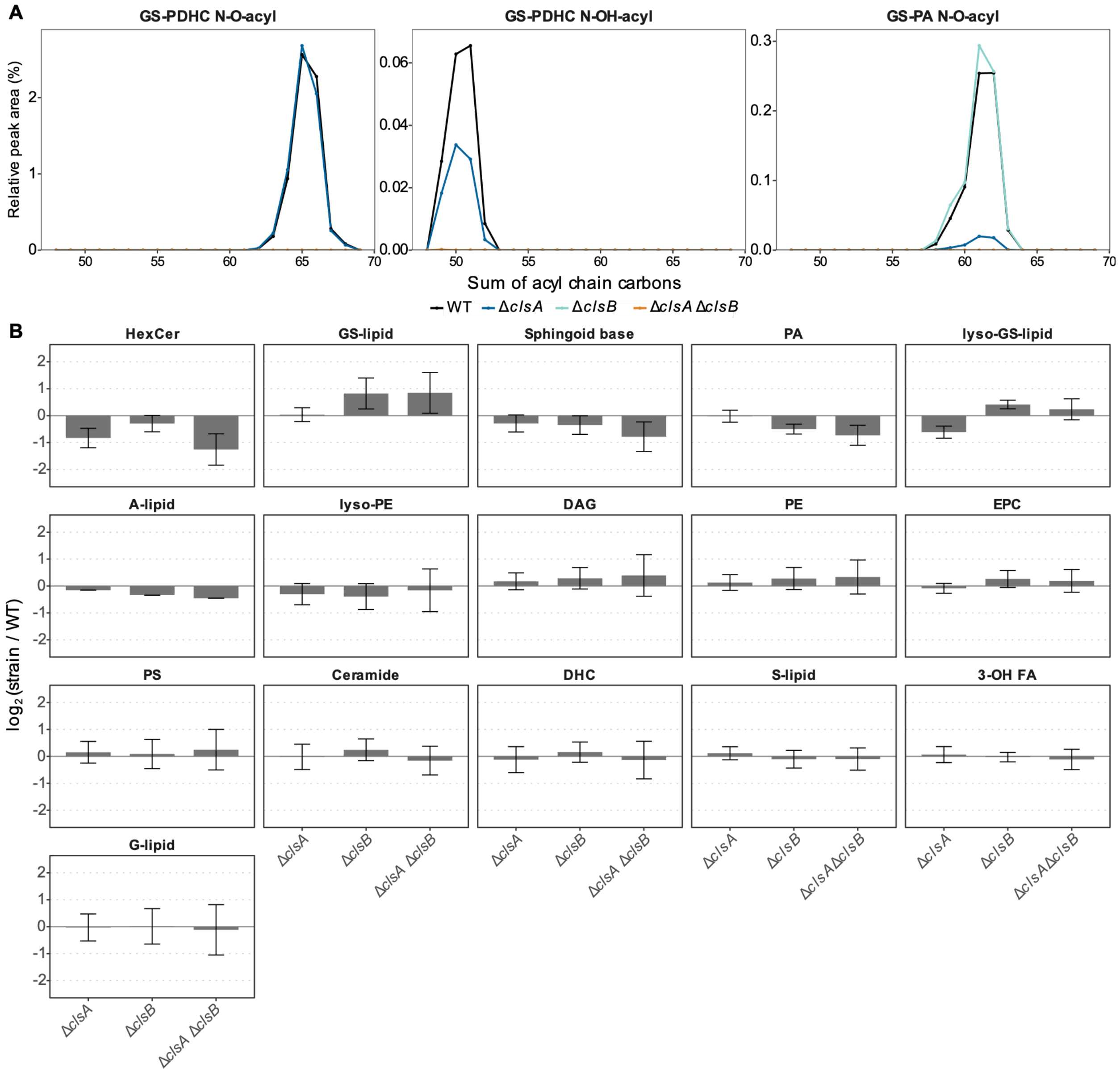


**Figure S7. Chain-length distributions of the glycyl-seryl classes across genotypes, and the lipid classes below the Fig. 7C threshold. A**. Chain-length distribution of each glycyl-seryl class (GS-PDHC N-O-acyl, GS-PDHC N-OH-acyl and GS-PA N-O-acyl) in each strain (key). Each class is plotted on its own y-axis. Where a class remained above 5% of the wild-type signal, its abundance-weighted mean chain length shifted by no more than 0.12 carbons. Loss of either gene therefore changes the amount of a class present without shifting its chain-length distribution. Negative ionization mode. **B.** Mean $\log_2$(fold change) relative to the wild type for every lipid class of the bacterial lipidome that does not meet the criterion applied in Fig. 7C. Bars are the mean across the molecules of a class detected in that genotype; error bars are the standard deviation of those per-molecule ratios and therefore show how uniformly a class responds. The number of molecules contributing to each bar is given in Table S3; A-lipid is a single molecule and has no error bar. All panels derive from three biological replicates per genotype. Lipid class abbreviations: PA, phosphatidic acid; PE, phosphatidylethanolamine; lyso-PE, monoacyl PE; PS, phosphatidylserine; DAG, diacylglycerol; Cer, ceramide; DHC, dihydroceramide; EPC, ceramide phosphoethanolamine; HexCer, hexosylceramide; sphingoid base, free (non-acylated) long-chain base; 3-OH FA, 3-hydroxy fatty acid; G-lipid, S-lipid and GS-lipid, glycine-, serine- and glycyl-seryl-conjugated lipids; lyso-GS-lipid, monoacyl GS-lipid; A-lipid, N-acyl amino-acid lipid with a $C_3H_7NO_2$ headgroup (alanine or an isomer), assigned from MS2; GS-PA, glycyl-seryl phosphatidic acid; GS-PDHC, glycyl-seryl phospho-dihydroceramide. N-O-acyl and N-OH-acyl denote the amide-linked subclasses. Molecule identities and class assignments are in Table S3.

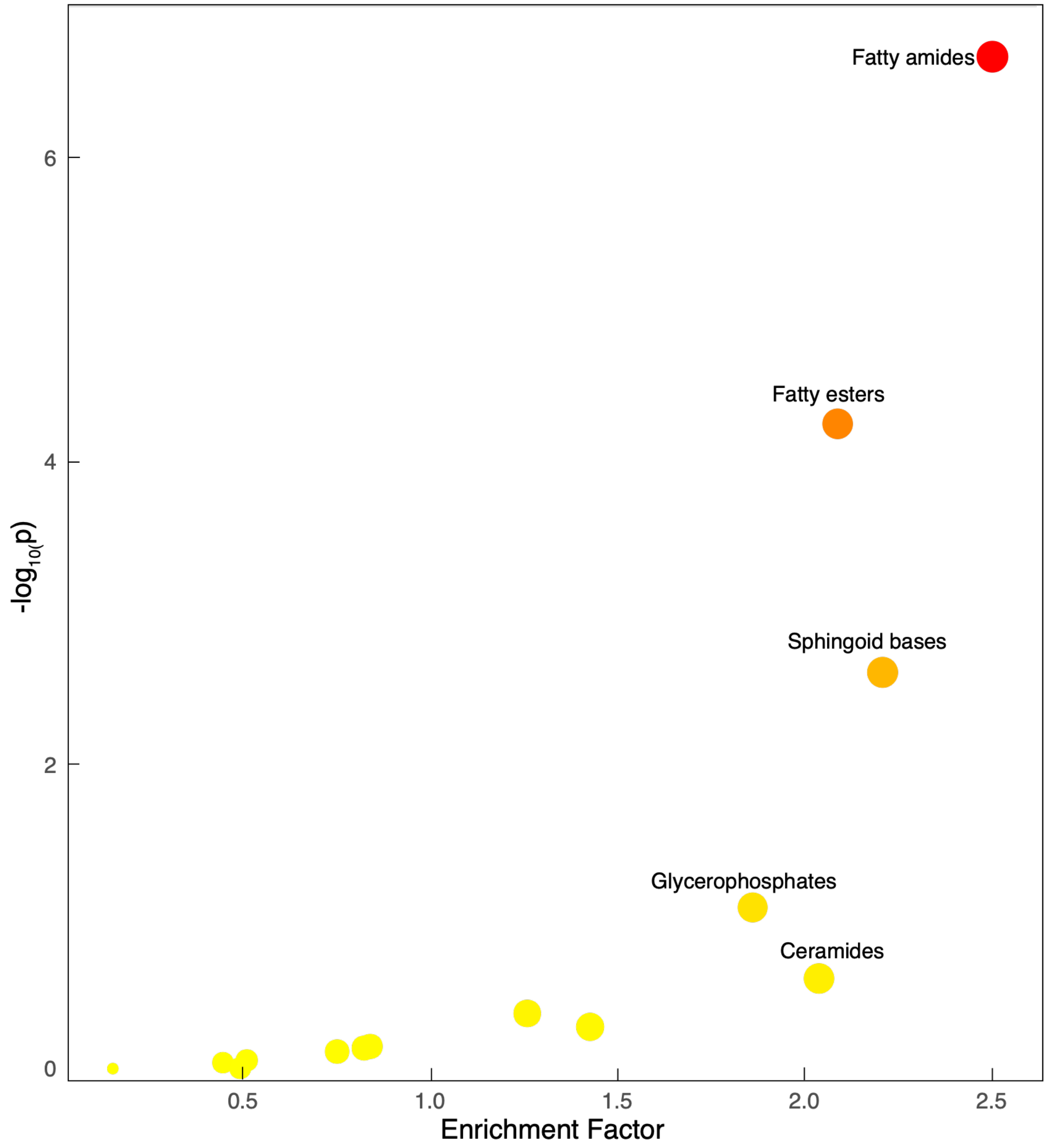


**Figure S8. Pathway enrichment analysis for main lipid categories.** Pathway analysis is performed with MetaboAnalyst 6.0, with default mummichog parameters with p-value cutoff of 0.0005 and lipid – main chemical class as the selected metabolite sets containing at least 3 entries.

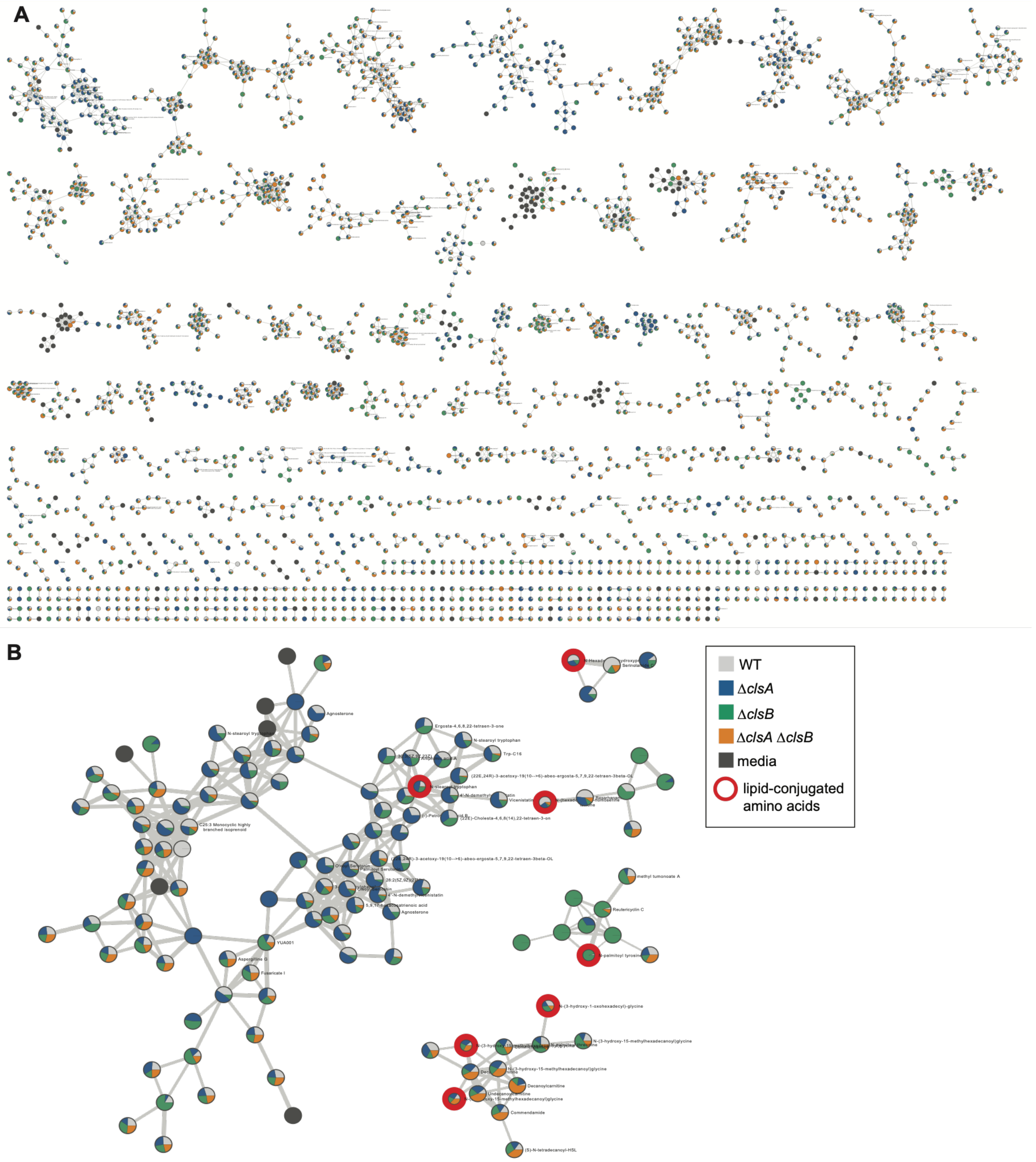


**Figure S9. Molecular networking shows chemically related metabolites in *B. fragilis* strains. A.** Feature-based molecular networking of all features (excluding singletons) from WT, Δ*clsA*, Δ*clsB*, Δ*clsA* Δ*clsB*, and extracted BHIS media. Each feature is a node, connected by an edge calculated from cosine similarity score. **B.** Subset of features connected by an edge to fatty amides highlighted in Fig. 8 (red border). Each pie chart in the node is colored by relative abundance of the feature in each strain.

**Table S1.** Primers and vectors used in this study.

**Table S2.** Metadata for genomics analysis.

**Table S3.** Lipidomics feature annotations, normalized abundance values, and cluster assignments.

**Table S4.** ICP-MS data.

**Table S5.** Metabolomics LC-MS data.

**Table S6.** Pathway Enrichment Data-Main Lipid.

**Table S7.** Pathway enrichment analysis for sub non-lipid categories.

### SUPPLEMENTAL METHODS

#### *Bacterial Growth and Strains*

*Strain generation*

All *B. fragilis* strains were grown in supplemented brain heart infusion (BHIS) medium composed of Bacto brain heart infusion (BD, Franklin Lakes, NJ), 1% (w/v) yeast extract (Fisher Scientific, Waltham, MA) and 0.1% w/v L-cysteine (Sigma-Aldrich, St. Louis, MS). Immediately before inoculation, filter sterilized 0.5% (w/v) hemin (Sigma-Aldrich, prepared in 0.1 N NaOH) and 0.1% (w/v) vitamin K1 (Sigma, prepared in 100% ethanol) was added, as we observed decreased bacterial viability if added before autoclaving or immediately after autoclaving (1). As BHIS medium ages, it supported progressively lower growth of *B. fragilis* str. P207 under our conditions; therefore, all experiments were conducted with refrigerated medium stored in the dark and prepared within the previous two weeks. *E. coli* strains were grown in Miller LB broth (BD) supplemented with filter-sterilized 300 μM diaminopimelic acid (Sigma-Aldrich) where appropriate for *E. coli* WM3064. Antibiotics were provided as 5 μg/mL erythromycin (*B. fragilis*) and 100 μg/mL carbenicillin (*E. coli*) for antibiotic vector selection and 100 ng/mL anhydrotetracycline (aTC) for inducible ssbfe1 counter-selection (1). For aTC-inducible complementation vectors, 50 ng/mL aTC was used due to the toxicity of this antibiotic to our *B. fragilis* strains (1). All growth was conducted at 37°C and all *B. fragilis* strains were grown in a Coy Anaerobic chamber in an atmosphere composed of 2-4% $H_2$ and 5% $CO_2$, with a balance of $N_2$.

Mutant strains were generated by allelic exchange using modified vectors described previously (1). Briefly, PCR fragments were amplified using KOD hot-start proof-reading polymerase (Novagen), digested with restriction enzymes (NEB), and ligated into the appropriate vector (listed below) using T4 DNA ligase (NEB) before transformation into chemically competent *E. coli* EC100D cells. Plasmid DNA was extracted from the positive colonies and verified by Sanger sequencing for the insert (RTSF Genomics Core at MSU, East Lansing, MI) or the whole plasmid (Plasmidsaurus, San Fransisco) depending on insert length. For conjugation into *B. fragilis* P207, constructs were transformed into chemically competent *E. coli* WM3064, a diaminopimelic acid (DAP) auxotroph. As described previously, *B. fragilis* P207 was back-diluted to very low OD (~0.005) and allowed to grow to a barely optically perceptible OD before combining with log-phase *E. coli* WM3064 (OD 0.3–0.8). Mixtures were spotted onto BHIS plates with 300 µM DAP (1). Mating spots were streaked for isolation on BHIS plates with 5 µg/mL erythromycin before counter-selection with 100 ng/mL aTC. Allelic exchange vectors (pLGB13-based) harbored a 1-kb homology region upstream and downstream of the gene(s) to be deleted with 10 remaining amino acids or fewer. The *clsA* mutant (lacking ptos_000612) was complemented using an aTC-inducible promoter, which is otherwise tightly repressed, due to suspected toxicity of the gene in *E. coli* (pMKS05-based) (1). The *clsB* mutant (lacking ptos_003252) was complemented with a constitutive *Bacteroides* promoter (*RpoD* sigma factor, bt_1311/BT_RS06635) (pMKS03-based) (2). Complementation constructs were integrated into the chromosome at the att site using an intN1 integrase (3). Primers, vectors, and strains are listed in Table S1.

#### *Cls conservation analysis*

*Alignment.* The *B. fragilis* P207 *clsA* and *clsB* genes, as well as *E. coli* K-12 MG1655 (NC_000913) *clsA*, *clsB*, and *clsC* genes, were translated and their encoded proteins aligned using default settings of the Geneious alignment algorithm (Geneious Prime 2024) (4, 5). Since ClsC failed to align with the other proteins, it was largely excluded from analysis. Proteins were aligned as a group as well as in pairs to generate the data presented in Fig. 1A–C.

*DeepTMHMM*. The ClsA and ClsB amino acid sequences from *B. fragilis* P207 and *E. coli* K-12 MG1655 were uploaded to DeepTMHMM (v. 1.0.42, https://dtu.biolib.com/DeepTMHMM) to predict transmembrane domains (6).

*Inter-Phylum Tree.* Representative taxa were selected from the top five phyla in the human gut (i.e., Bacteroidota, Bacillota, Actinomycetota, Fusobacteriota, and Pseudomonadota) (metadata listed in Table S2) (7, 8). Even though not generally present in the human gut, one taxon from the class Alphaproteobacteria (*Sphingomonas paucimobilis*) was also added as a comparison to the other Pseudomonadota classes. Genome sequences were downloaded from NCBI, annotated *cls* genes were translated and their encoded proteins aligned in Geneious. One genome per species was used, aside from the two *B. fragilis* strains, to compare the diversity of *cls* genes within particular organisms (i.e., *Turicibacter sanguinis* has five annotated *cls* genes). The phylogenetic tree was reconstructed using the neighbor-joining method via Geneious (100 bootstrap replicates) and then visualized using R (see below). No outgroup was specified during tree reconstruction.

*Intra-*Bacteroides *Tree.* The TIGR04265 domains in *B. fragilis* ClsA and ClsB were identified using the conserved domain database of NCBI (9). Proteins containing this domain were identified by querying the Interpro database and the amino acid sequences from all *Bacteroides* matching entries were downloaded, excluding uncultured strains and taxa without an assigned species name. Alignment, tree reconstruction, and tree visualization were performed as with the inter-phylum tree, except that *E. coli* K-12 MG1655 ClsA was specified as an outgroup.

*Genomic Neighborhoods.* WebFlags2 was used to query genomic neighborhoods around *B. fragilis* P207 *clsA* and *clsB* genes (10). The top 200 hits from the search algorithm results were saved and then manually curated for brevity by removing extraneous/redundant neighborhoods.

#### *Growth curve assays*

*B. fragilis* strains were grown overnight anaerobically at 37°C with antibiotic selection (i.e., erythromycin) and without inducer. In the morning, stationary phase cultures were back-diluted to an optical density at 600 nm ($OD_{600}$) of 0.05 and allowed to grow to early log phase ($OD_{600}$ ~0.3) in the presence of aTC inducer. Each strain was then diluted again to 0.025–0.05 for growth curves in a 96-well plate format. Cells were grown in technical triplicate in non-tissue culture–treated polystyrene plates (GenClone, El Cajon, CA) with BreatheEasy sealing membranes (Sigma-Aldrich), which are translucent, gas-permeable membranes that individually seal each well and minimize condensation. $OD_{600}$ was measured with a Tecan Infinite M Nano plate reader (Tecan, Männedorf, Switzerland) for 24 h every 10 min with 30 s of shaking and 10 s of settling before each read. Three biological replicates were performed for each growth condition.

To expose cells to stress conditions, BHIS medium was prepared as above but modified as below. For pH assays, cells were pelleted at 16,000 × *g* for 1 min and then resuspended in aTC-containing BHIS medium adjusted to the appropriate pH using HCl or NaOH with an Orion Star pH probe (ThermoFisher). For deoxycholate and SDS assays, 0.01% (w/v) deoxycholate sodium salt (Sigma) or 0.002% (w/v) SDS (Lab Scientific, Danvers, MA) was added to cultures from filter-sterilized 1% (w/v) stocks dissolved in UltraPure water. For $Na^+$ and $K^+$ assays, NaCl (Fisher Scientific) and KCl (Fisher Scientific) were dissolved to a 4× concentration of each cation in BHIS medium and then filter sterilized before being added to cells to yield the final concentration of cells and cation (i.e., 300 mM). To determine how much additional NaCl or KCl to add, the average $Na^+$ and $K^+$ content of our BHIS medium was determined using ICP-MS (see below for methods), yielding 130 mM for $Na^+$ and 33 mM for $K^+$ (Table S4). For ionophore assays, 2 µL of a concentrated ionophore stock (monensin A or nigericin; Sigma-Aldrich) prepared and diluted in ethanol was added to cultures to yield final concentrations of 0.8 µg/mL monensin A or 0.1 µg/mL nigericin. An equivalent volume of ethanol without any ionophore was added to non-treated conditions during these experiments as a vehicle control.

To calculate maximum $OD_{600}$ and doubling time, the time of maximum growth rate was identified by fitting a $4^{th}$-degree polynomial to each biological replicate with the R package ipolygrowth, which uses ordinary least squares estimation (11). The growth data were then trimmed to 6 h before and after this time point, and then the polynomial was refit to improve metric estimation across diverse growth curves. Per ipolygrowth, peak growth rate is defined as the first derivative of the slope at peak growth time, and the doubling time is ln(2)/peak growth rate. The max $OD_{600}$ value was determined by finding the point at which the growth curve reached an asymptote. In Figure S3, $\log_2$(fold change) compares these metrics from each strain replicate to the average of biological replicates from the wild type.

#### Microscopy

*B. fragilis* P207 cell cultures were grown overnight under antibiotic selection and then back-diluted into medium containing inducer (aTC) without antibiotic selection to reach log-phase growth. All cultures were removed from the 37ºC incubator but were maintained under anaerobic conditions, with only two or three cultures removed for microscopy at a time. $OD_{600}$ was measured and 1 µL of culture was placed onto a 1% (w/v) agarose pad made of BHIS medium as described (12). Cells were visualized by phase-contrast microscopy on a Leica DMI6000 B inverted microscope with a Hamamatsu ORCA-R2 10600 camera with a 63× Plan Apo objective. Each strain was visualized on four different days to generate four biological replicates. The two images per replicate from each strain/day were chosen for analysis.

Chosen images were processed using the MicrobeJ (v. 5.13l) plugin cell size analysis software in Fiji (v. 1.54k) at −5 sensitivity and excluding cells below 0.5 µm in length, since our experience demonstrated the vast majority of these calls were image artifacts (13). Called cells were then manually curated by removing cells that were overlapping or out of focus. The resulting cell lengths and widths were exported and analyzed in R. Cell volume was calculated according to the formula $V = [(w^2 \times \pi/4) \times (l - w)] + (\pi \times w^3/6)$, where $w$ is cell width and $l$ is length, and Epanechnikov kernel density estimates of the cell data were generated to visualize broader trends (14, 15). To test for differences between strains, a mixed-effect regression model [$\log_{10}$(cell width) ~ strain + 1|biological replicate] was used for differences in means; for differences in data distribution (i.e., variance from the mean), an *F*-test was used.

#### *Gene expression assays*

*Stress condition exposure*: Strains were grown overnight and back-diluted as described for the growth assays. For comparison of *cls* gene expression at log and stationary phases, *B. fragilis* WT, Δ*clsA* and Δ*clsB* strain cultures were again diluted to an $OD_{600}$ of 0.05 without inducer after back-diluted cultures reached log phase ($OD_{600}$ ~0.3). Two hours later, samples were collected and cells were pelleted at 16,000 × *g* for 1 min and resuspended in 1 mL of Trizol (Invitrogen). Stationary-phase samples were collected at 24 h. Samples were randomized to minimize collection bias across replicates. Samples were then frozen at −80°C until RNA extraction. For acute toxicity assays, back-diluted WT *B. fragilis* cultures that reached $OD_{600}$ ~0.3–0.5 were exposed to a specified condition (i.e., pH, SDS, deoxycholate, $K^+$, monensin A, or nigericin, as described for the growth assays) aerobically in a 1.5-mL Eppendorf tube and incubated for 20 min. Cells were pelleted by centrifugation at 17,000 × *g* for 1 min in the same tubes and resuspended in 1 mL Trizol before being frozen at −80°C.

*RNA extraction*: RNA was extracted using a phenol-chloroform method as described previously with modifications (4). Trizol samples were thawed and heated at 65ºC for 10 min. Then, 200 µL of chloroform (Macron Fine Chemicals) was added and samples were thoroughly vortexed. After a 5-min incubation at room temperature, samples were centrifuged at 17,000 × *g* for 15 min at 4°C. The upper, aqueous phase was transferred to a fresh RNase-free microcentrifuge tube. A 0.7× volume of RNA-grade 100% isopropanol (Fisher) was added and then samples were frozen at −80°C for RNA precipitation. Following this freezing step, tubes were centrifuged at 17,000 × *g* at 4°C for 30 min to pellet RNA. Pellets were washed in RNA-grade 70% (v/v) ethanol twice, air-dried on the bench for 10 min, and then suspended in 50 µL RNase-free $H_2O$. An RNeasy Extraction Kit (Qiagen) was used to treat our samples with Turbo DNase (Invitrogen) using the standard protocol. Briefly, DNase I was added to the silica membrane containing RNA. RNA was then eluted in 50 µL RNase-free water and stored at −80°C until RT-qPCR.

*RT-qPCR*: Samples were run in technical triplicates in non-transparent PCR plates with optically clear sealing film (Applied Biosystems). *Ct* values were determined using a QuantStudio5 (Applied Biosystems) instrument and a Luna One-Step RT-qPCR kit (NEB) for target genes in each sample (4). Using *B. fragilis dnaN* (ptos_002600, encoding DNA polymerase subunit beta) as a normalization gene since it did not fluctuate previously in deoxycholate stress experiments, the − ΔΔCt and fold change values for *clsA* and *clsB* transcripts were calculated using the formulas $\Delta\Delta Ct = [(Ct_x - Ct_{\mathrm{ref}})_{\mathrm{test}} - (Ct_x - Ct_{\mathrm{ref}})_{\mathrm{control}}]$ and $\text{fold change} = 2^{-\Delta\Delta Ct}$.

#### *Inductively coupled plasma mass spectrometry*

Cells were grown overnight and back-diluted to place them into log-phase growth as described in the growth assays. At $OD_{600}$ 0.3-0.4, cells were removed from the anaerobic chamber for sample collection. If cells were exposed to a stress, they were incubated for 20 min aerobically at room temperature with either 0.01% (w/v) deoxycholate, 0.8 µg/mL monensin A or 0.1 µg/mL nigericin. After taking an aliquot for counting colony-forming units on BHIS agar, 5 mL of culture were transferred to a pre-weighed 15 mL metal-free tube (LabCon, Petaluma CA). The culture was then spiked with gadolinium-DOTA (Gd-DOTA, Macrocyclics, Cat # M-147) prepared in UltraPure water to a concentration of 40 µM. Washing cells, even with the BHIS medium cells were grown in, perturbs intracellular concentrations of small alkali metals like $Na^+$ and $K^+$, as well as numerous other elements, making their quantification difficult (Fig. S5A). Using a Gd-DOTA spike enabled us to remove any washing steps in the ICP-MS sample preparation process since it is a membrane impermeable complex and therefore primarily resides in the extracellular space (16, 17).

For the wash experiment, Gd-DOTA was spiked into cell cultures to 40 µM. For wash conditions, 5 mL of cells were washed twice by pelleting cells at 7,100 $\times$ g for 10 min, the supernatant was removed, and the cells were then resuspended in either 25 mL of 250 mM sucrose or BHIS media. Cells were pelleted one final time at 7,100 $\times$ g for 10 min before resuspending the pellet in 5 mL of wash solution. Then, 5 mL of unwashed cells (i.e., no previous centrifugation) and resuspended washed cells were transferred to a metal-free tube where they were pelleted at 4,000 $\times$ g for 10 min. A 400 µL of the supernatant was transferred to a fresh metal-free tube for analysis and the remainder of the supernatant from the 5 mL culture was removed in two steps: most of the supernatant was removed via aspiration, then the cells were centrifuged for 1 min at 4,000 $\times$ g to remove drips of supernatant from the sides of the tube, then the rest of the supernatant was removed.

For *cls* and treatment comparison experiments, 5 mL of unwashed cells were pelleted for 3 min at 7,000 $\times$ g after which 0.5 mL of supernatant was transferred to another pre-weighed 15-mL conical tube for analysis. The remainder of the supernatant was removed carefully as above but spun at 7,000 $\times$ g. Cell pellets and supernatants were stored at −20 $^{o}$C until acid digestion.

Weighed cell pellets and supernatant were dried in an oven at 70 $^{o}$C overnight, then digested using 70% $HNO_3$ acid at 70 $^{o}$C overnight. The resulting digested solution were then diluted with ultrapure deionized water to 3% nitric acid. After weighing the resulting solutions, then the further diluted (50x) solutions were also prepared with 3% nitric acid to measure highly abundant elements such as P, S, and K by ICP-MS. All standards, blanks, and ICP-MS samples were prepared using ultra-trace metal grade nitric acid (70%, Fisher chemical, Cat# A467-250), ultrapure water (18.2 MΩ·cm at 25 °C) obtained from a Milli-Q IQ 7000 system (Millipore Inc., Billerica, MA), metal free polypropylene conical tubes (15 and 50 mL, Labcon, Petaluma, CA, USA), and trace metal grade pipette tips (Labcon, Petaluma, CA, USA). All solutions were weighed for accurate elemental determination using the XSR205 DU semi-analytical balance (Mettler Toledo, Columbus, OH, USA).

The completed ICP-MS samples were analyzed using an Agilent 8900 Triple Quadrupole ICP-MS (Agilent Technologies, Santa Clara, CA, USA) equipped with the Agilent SPS 4 Autosampler, integrated sample introduction system (ISiS), x-lens, and MicroMist nebulizer. Daily tuning of the instrument was accomplished using a manufacturer-supplied tuning solution containing Li, Co, Y, Ce, and Tl. Global tune optimization was based on optimizing intensities for $^{7}$Li, $^{89}$Y, and $^{205}$Tl while minimizing oxides ($^{140}Ce^{16}O^+/^{140}Ce^+ < 1.5\%$) and doubly charged species ($^{140}Ce^{++}/^{140}Ce^+ < 2\%$). Following global instrument tuning, gas mode tuning was accomplished using the same manufacturer supplied tuning solution in both KED and Oxygen mode (using 100% UHP He and $O_2$ gas, Airgas) at 5 mL/min. Specifically, intensities for $^{59}Co^+$, $^{89}Y^+$, and $^{205}Tl^+$ were maximized while minimizing oxides ($^{140}Ce^{16}O^+/^{140}Ce^+ < 0.5\%$) and doubly charged species ($^{140}Ce^{++}/^{140}Ce^+ < 1.5\%$) with short term RSDs < 3.5%.

To prepare ICP-MS standards, the multi element standard IV-65024 (Inorganic Ventures, Christiansburg, VA, USA) that contains 10 ppm As, Ca, Cd, Co, Cr, Cu, Fe, Gd, K, Mg, Mn, Mo, Na, Ni, P, Pt, S, Se, V, and Zn, was diluted with 3% (v/v) nitric acid in ultrapure water to a final element concentration of 0 (blank), 0.01, 0.05, 0.1, 0.5, 1, 3, 5, 10, 15, 20, 40, 100, 200, and 400 ppb. Internal standardization was accomplished inline using the ISIS valve and a 200 ppb internal standard solution in 3% (v/v) nitric acid in ultrapure water consisting of Bi, In, $^{6}$Li, Sc, Tb, and Y (IV-ICPMS-71D, Inorganic Ventures, Christiansburg, VA, USA). The isotopes selected for analysis were $^{23}$Na, $^{31}$P, $^{32}$S, $^{39}$K and $^{157}$Gd with $^{45}$Sc, $^{89}$Y, $^{115}$In, and $^{159}$Tb used for internal standardization. For accurate quantification of $^{31}$P and $^{32}$S, these isotopes were measured in the Oxygen mode.

To determine intracellular element counts, the Gd ratio was calculated according to the formula $\mathrm{Gd_{ratio}} = [\mathrm{Gd}]_{\mathrm{cells+residual\ media}}/[\mathrm{Gd}]_{\mathrm{supernatant}}$. Since Gd-DOTA is membrane-impermeable, most of the Gd signal will be from elements in the residual medium and not from intracellular content (16, 17). By calculating the Gd ratio, the amount of residual supernatant-derived elements can be calculated as $[\mathrm{Elements}]_{\mathrm{residual\ media}} = [\mathrm{Elements}]_{\mathrm{supernatant}} \times \mathrm{Gd_{ratio}}$ and then subtracted from the counts for the digested cell pellet to obtain only intracellular elemental counts (i.e., $[\mathrm{Elements}]_{\mathrm{intracellular}} = [\mathrm{Elements}]_{\mathrm{cell+residual\ media}} - [\mathrm{Elements}]_{\mathrm{residual\ media}}$).

The number of atoms per cell was derived by normalizing to the number of colony-forming units for each sample. Due to variability in colony-forming unit determination, at least two aliquots were collected for each individual culture, and each was serially diluted. Then, four replicate 5-µL spots were added to plates and colonies were counted after 18 h of anaerobic incubation at 37°C. To quantify intracellular elemental concentration, the average strain-specific cell volume was determined and then the molar concentration calculated for each element from the number of atoms per cell. The ΔmM concentrations were calculated by subtracting the concentration of an element in the medium from that in the intracellular space.

#### *Lipidomics assays*

WT, Δ*clsA*, Δ*clsB*, and Δ*clsA* Δ*clsB* strains were grown overnight to stationary phase ($OD_{600}$ 1.4) and then back-diluted to $OD_{600}$ 0.05 as described for the growth assays. Then, 1.5 mL of culture at an $OD_{600}$ of 0.7 was pelleted at 16,000 × g for 5 min and then submitted on dry ice to the University of Tennessee-Knoxville Biological and Small Molecule Mass Spectrometry Core (BSMMSC; RRID: SCR_021368) for targeted lipidomic analysis of membrane lipids, including ceramides, sphingolipids, phospholipids and cardiolipins. Lipids were extracted as previously described (18, 19). Briefly, bacterial cells pellets were resuspended in 1 mL of extraction solvent (15: 15: 5: 1: 0.18 95% ethanol: water: diethyl ether: pyridine: 4.2 N ammonium hydroxide) and glass beads (150-212 µm) were added. The resuspended cells were incubated at 60 ºC for 20 min and then centrifuged at 16,200 × g for 10 min. The supernatant was collected and this extraction process repeated with an additional 1 mL of extraction solvent. The combined supernatant was dried under a steady stream of nitrogen. Prior to mass analysis the dried extracts were resuspended in 9:1 methanol: chloroform. The lipidomic analyses were performed using a previously validated method (19) on an ultra high performance liquid chromatography system coupled to a high resolution mass spectrometer (UHPLC-HRMS). The chromatographic separations were carried out using Vanquish Horizon LC system (Thermo Scientific) and reversed phase separations as described previously (19). The eluent was introduced to an Exploris 120 mass spectrometer (Thermo Scientific) via electrospray ionization for high resolution analysis. Following the analysis, the lipids were annotated and the peak area integrated using MS-DIAL (20-23). Annotation was performed separately for positive- and negative-ion-mode data using an MS1 tolerance of 0.01 Da, an MS/MS tolerance of 0.025 Da, and a 2-min retention-time tolerance. Candidate features were filtered against blank samples, and features with incorrect exact-mass assignments or ghost peaks were removed, including through the MS-DIAL/MS-CleanR workflow applied after initial annotation. Because many lipid features lacked MS/MS spectra, annotations were manually reviewed, anchoring on the masses reported by Franfater et al. {Frankfater, 2023 #83}: retained features were required to fall within 5 ppm of the expected exact mass, MS/MS spectral matches were manually inspected when available, and peaks were retained only when the reported signal-to-noise ratio was ≥3. Retained lipid assignments should therefore be considered annotated LC-MS features supported by accurate mass, retention behavior, blank filtering, and manual review, with MS/MS support where available. The peak areas were then used for further normalization and analysis. Lipid abundances were normalized across samples to $OD_{600}$ of sample cultures.

Peak area for a metabolite was first normalized to the optical density of a sample, then relative lipid abundance was determined by normalizing to the total peak area of all lipids detected in each UHPLC−MS/HRMS mode (i.e., positive versus negative). Particular ionization modes were prioritized for detailed analysis of lipids better detected with that charge. Principal component analysis was performed on the normalized lipid-abundance (peak-area) data as an unsupervised overview of whole-lipidome differences among the WT and cls mutant strains, with minimum volume-enclosing ellipses estimated using Khachiyan's algorithm.

To account for feature-specific differences in MS response, a follow-up UHPLC-MS/HRMS run analyzed biological triplicates of all four strains alongside a seven-point pooled-extract serial dilution series spanning six orders of magnitude (1× to $10^6$× dilution, in triplicate). Most lipid features were detected only at the two or three least-diluted levels (1×, 10×, and 100×); a feature was retained for quantification when it was detected at a minimum of two consecutive dilution levels in at least two of three replicates per level. For each retained feature, a calibration curve was fit by linear regression of integrated peak area against relative lipid amount, defined as the reciprocal of the dilution factor (1/dilution). For each experimental sample, a calibration-derived "amount equivalent" was calculated for each lipid feature as (area − intercept)/slope and expressed in units of the original pool concentration; these amount equivalents were further normalized to the mean amount equivalent in WT samples to express each lipid in the *cls* deletion strains as a fold-change relative to WT. Calibration parameters and per-feature quantification-confidence flags (including two-point fits, extrapolated values, near-detection-limit pools, and negative slopes) are provided in Table S3. k-means clustering of this calibrated dataset was used as an exploratory approach to group lipid features by their abundance patterns across strains; cluster 1 (km1) is *clsB*-associated, cluster 3 (km3) is *clsA*-associated, and cluster 2 (km2) comprises the remaining features, with per-feature cluster assignments provided in the km_cluster column of Table S3 and the clustering shown in Fig. 6D. The OD-normalized dataset and the dilution-series calibration dataset derive from independent UHPLC-MS/HRMS runs and are not directly comparable feature-by-feature, so within-run comparisons were used throughout. Because authentic standards were not available for all annotated lipid classes, these values were treated as semi-quantitative rather than absolute measurements of lipid abundance.

#### ***Metabolomics assays***

WT, Δ*clsA*, Δ*clsB*, and Δ*clsA* Δ*clsB* strains were grown in 50mL BHIS media for 24 hr. Each culture was then sonicated for 60 s and extracted with equal volume of ethyl acetate three times. The ethyl acetate layer was collected after each extraction and dried using rotary evaporation, transferred to vials, and stored at -80°C. All solvents were of high-performance liquid chromatography (HPLC) grade and purchased from Sigma Aldrich.

Extracts were prepared for metabolomics analyses at 1 mg/mL in 50% methanol (MeOH). Data was acquired using a Bruker timsTOF Pro2 (Bruker-Daltonics, Billerica, MA, USA) coupled to an Agilent 1290 Infinity II Bio UHPLC (Agilent, Santa Clara, CA, USA). Each sample (2 uL) was injected in technical triplicate at random using an Acquity UPLC HSST3 column (2.1 x 150mm, 1.8 $\mu$M). Samples were eluted using a 0.3 mL/min gradient of mobile phases A (0.1% formic acid in water) and B (0.1% formic acid in acetonitrile) employing the following conditions: 1 min hold at 5% B, 1 min ramp to 15% B, 6 min ramp to 100% B, 3 min hold at 100% B, 0.1 min ramp back to 5% B and a re-equilibration hold at 5% B for 1.4 min.

Data acquisition was performed in positive ionization mode using an ESI source, with a collision energy of 10eV, capillary voltage of 4500 V, dry temperature of 22°C, sheath gas temperature of 22°C, mass range of 50-2000 m/z, and mobility (1/Ko) range of 0.55-1.90 V.s/cm2. Fragmentation data were acquired with a collision energy of 50 eV, with 2 PASEF MS/MS scans per cycle, for a total cycle of 0.53 s.

Once acquired, MS data were preprocessed using Bruker MetaboScape® version 9.0.1 (Bruker-Daltonics, Billerica, MA, USA) using the MCube T-Rex 4D Metabolomics workflow for peak picking and alignment. The intensity threshold was experimentally determined by comparison of the baseline noise from samples and 50% methanol (MeOH) blanks resulting in an intensity threshold of 1500 counts. The resulting feature table was then processed using mpactR (24) with the following parameters: mispicked peak correction - ringing mass window of 0.5 atomic mass units (AMUs), isotopic mass window of 0.01 AMU with a maximum isotopic mass shift of 3 AMUs, and a $t_R$ window of 0.05; in-source ion filtering threshold of 0.95 Spearman correlation; median coefficient of variation (CV) of technical replicates of 0.5; and blank filtering using 50% MeOH blanks at a 0.05 threshold.

*In-silico* formula prediction was performed with MetaboScape® 2024b. Small molecules and lipid databases were used for annotations with a 5 ppm cutoff. Small molecule databases included: NPAtlas (25), MS-DIAL tandem mass spectral standards (23, 26), Bruker MetaboBASE Personal Library 3.0, Bruker NIST mass MSMS spectral library. Lipid databases included: Bruker rule-based lipids annotation and Lipid Maps database (27). Annotations were further verified by comparing fragmentation patterns using public data, when available, or using the Competitive Fragmentation Modeling for Metabolite Identification (CFM-ID) spectra prediction (26).

Peak intensity for a metabolite was normalized to the total peak intensity of all features detected in the sample. Molecular networking was generated as previously described (28) in GNPS2: Global Natural Products Social Molecular Networking 2 (http://gnps2.org/). A minimum cosine similarity score of 0.7 with at least six matching peaks, parent mass tolerance of 2 Da, and fragment ion tolerance 0.5 Da were selected to generate consensus spectra. Files were imported into CytoScape v(3.10.3) and nodes were arranged with yFiles organic layout plugin (29).

Pathway analysis was performed with MetaboAnalyst 6.0 functional analysis [LC-MS] module (30). A formatted peak intensity table was uploaded to the web server (https://metaboanalyst.ca), then normalized by sum and log transformed. Default setting of 5.0 ppm mass tolerance, mummichog algorithm v2.0 with p-value cutoff of 0.0005, against the lipids – main chemical class, and non-lipids – sub chemical class pathway libraries with at least 3 entries in each pathway. Fold change analyses for metabolites from enriched pathways were performed by dividing the average value of the metabolite in mutant strains by the average value in the wildtype strain, and results were plotted in R v(4.5.0).

#### *Data analysis and reproducibility*

All data analyses were performed using base R (v4.3.3) (31) as well as these packages and resources for the following analysis: 1) data visualization: ggplot2 (32), a colorblind safe palette (33), colorBlindness (34), RColorBrewer (35), ggbiplot (36), ggforce (37), circlize (38), ComplexHeatmap (39), mdthemes (40), ggtext (41); 2) data manipulation: readxl (42), tidyr (43), matrixStats (44), data.table (45), magrittr (46), scales (47), dplyr, tidyverse (43); 3) tree reconstruction: treeio (48) and ggtree (49, 50) ipolygrowth (11) and 4) statistics: lme4 (51, 52), ggubr (53) and glmnet (54, 55). Code used for all data analysis and figure generation, except for the WebFlags neighborhoods, is published on GitHub (https://github.com/mschnizlein/bfrag_cardiolipin). Original datasets were also posted on GitHub. Lipidomics data alongside appropriate metadata were deposited on MetaboLights (MTBLS11891 www.ebi.ac.uk/metabolights/MTBLS11891). Untargeted metabolomics were deposited to MassIVE (https://massive.ucsd.edu) (deposit in progress at time of submission).
